# RASSF1C oncogene elicits amoeboid invasion, cancer stemness and invasive EVs via a novel SRC/Rho axis

**DOI:** 10.1101/2021.01.05.425393

**Authors:** Maria Laura Tognoli, Nikola Vlahov, Sander Steenbeek, Anna M. Grawenda, Michael Eyres, David Cano-Rodriguez, Simon Scrace, Christiana Kartsonaki, Alex von Kriegsheim, Eduard Willms, Matthew J. Wood, Marianne G. Rots, Jacco van Rheenen, Eric O’Neill, Daniela Pankova

**Affiliations:** CRUK/MRC Oxford Institute for Radiation Oncology, ORCRB, Oxford, OX3 7DQ, UK; Department of Oncology, ORCRB, Oxford, OX3 7DQ, UK; Molecular Pathology, Oncode Institute, The Netherlands Cancer Institute, Amsterdam, The Netherlands; University of Groningen, University Medical Center Groningen, Groningen, The Netherlands; Cancer Research UK Edinburgh Centre, MRC Institute of Genetics & Molecular Medicine, The University of Edinburgh, Western General Hospital, Crewe Road South, Edinburgh, EH4 2XR, UK; Department of Physiology, Anatomy and Genetics, University of Oxford, Oxford, UK & La Trobe Institute for Molecular Science, La Trobe University, Melbourne, Australia; Department of Paediatrics, University of Oxford, Oxford, UK

## Abstract

Cell plasticity is a crucial hallmark leading to cancer metastasis. Upregulation of Rho/ROCK pathway drives actomyosin contractility, protrusive forces and contributes to the occurrence of highly invasive amoeboid cells in tumors. Cancer stem cells are similarly associated with metastasis, but how these populations arise in tumors is not fully understood. Here we show that the novel oncogene RASSF1C drives mesenchymal to amoeboid transition and stem cell attributes in breast cancer cells. Mechanistically, RASSF1C activates Rho/ROCK via SRC mediated RhoGDI inhibition, resulting in generation of actomyosin contractility. Moreover, we demonstrate that amoeboid cells display the cancer stem cell markers CD133, ALDH1 and the pluripotent marker Nanog; are accompanied by higher invasive potential *in vitro* and *in vivo;* and employ extracellular vesicles to transfer the invasive phenotype to target cells and tissue. Importantly, the underlying RASSF1C driven biological processes concur to explain clinical data: namely, methylation of the RASSF1C promoter correlates with better survival in early stage breast cancer patients. Therefore, we propose the use of *RASSF1* gene promoter methylation status as a biomarker for patient stratification.

## Introduction

Cell invasion and migration are essential processes during cancer progression and metastatic dissemination. In different tissue contexts, cancer cells use distinct modes of invasion, moving either as collective groups or single cells, adopting mesenchymal or amoeboid motility ^1,2^. Mesenchymally invading single cells use integrins and metalloprotease-dependent degradation of the extracellular matrix to invade and metastasize to distant organs. Downregulation of integrins or inhibition of metalloproteases results in mesenchymal-amoeboid transition (MAT) and adoption of amoeboid type of invasion. Amoeboid cancer cells are characterized by upregulation of Rho/ROCK signaling pathway, which drives reorganization of the actin cytoskeleton and leads to generation of protrusive forces. Enhanced Rho/ROCK signaling promotes MLCK mediated phosphorylation of myosin light chain 2 (pMLCII), which in turn induces reduction of stress fibers and formation of cortical actomyosin ^3,4^. These major changes in actin architecture enable cancer cells to remodel components of the extracellular matrix, and to squeeze into the surrounding tissue. Interestingly, invading cells can employ a hybrid mode of motility, switching between mesenchymal-amoeboid (MAT) and amoeboid-mesenchymal transition (AMT), depending on the tissue specificity of the matrices ^5,6^. This cellular plasticity allows cancer cells to initiate a lesion and adopt appropriate invasive modes for each step of the metastatic process. Cancer dissemination is a complex mechanism, and the exact sequence of molecular events that lead to metastatic colonization of distant sites of the body is not well understood. Extracellular vesicles (EVs) have recently been demonstrated to influence epithelial-mesenchymal transition (EMT) and migration in both an autocrine and non-cell autonomous manner ^7,8^. Extracellular vesicles are small lipid bilayer vesicles released from internal compartments such as multivesicular bodies (exosomes), or via blebbing from the plasma membrane (ectosomes, microvesicles) and have been implicated in promoting metastasis ^9,10^. Extensive membrane blebbing facilitated by Rho-driven motility has been associated with increased shedding of vesicles ^11–13^. It has recently been reported that RhoA-driven tumor progression is attributed to loss of the Hippo pathway scaffold and tumor suppressor RASSF1A ^14^. Epigenetic suppression of the *RASSF1-1α* promoter has been associated with poor cancer survival ^15,16^ due to suppression of RASSF1A transcript expression. However, this event also represents an epigenetic switch and expression of an alternative isoform, RASSF1C, from the internal *RASSF1-2γ* promoter ^17^. Although methylation of the *RASSF1* gene has been investigated in a large number of clinical studies and has been considered as a potential biomarker for breast cancer progression ^18^, how methylation of *RASSF1-2γ* influences disease outcome has not been addressed. RASSF1C has been shown to activate SRC kinase and induce an invasive phenotype, both *in vitro* and *in vivo*^17,19^. SRC plays a crucial role in cancer cell plasticity and has been described as a key player in epithelial-mesenchymal transition (EMT) in solid tumors, via its association with FAK and β-integrins ^20^. SRC can also regulate small Rho-GTPases by phosphorylation of RhoGDI ^21^. Here we identified a novel mechanism where SRC promotes amoeboid invasiveness via RASSF1C-SRC-RhoGDI signaling and Rho/ROCK/pMLCII activation. We show that SRC activation by RASSF1C leads to inhibitory phosphorylation of RhoGDI, followed by RhoA/ROCK upregulation, which in turn drives mesenchymal-amoeboid transition (MAT). We also demonstrate that molecular events driven by MAT are associated with cancer stemness and that amoeboid cells express the cancer stem cell markers ALDH1, CD133 and the pluripotency marker Nanog. Additionally, our data illustrate that RASSF1C-SRC-Rho activity results in release of EVs that transfer stemness, invasive ability and metastatic behavior to recipient cells *in vitro* and *in vivo*.

Detailed analysis of the *RASSF1* promoter *(RASSF1-1α* vs *RASSF1-2γ)* in early stage breast cancer supports the association of *RASSF1C* with adverse prognosis and offers an optimized biomarker for patients.

## Results

### The RASSF1C oncogene promotes mesenchymal-amoeboid transition

RASSF1C oncogene is one of the two main isoforms encoded by the *RASSF1* gene (Supp. Fig. 1A). The major isoform, RASSF1A, is a *bona fide* tumor suppressor epigenetically inactivated in numerous cancers (lung and breast cancer, among others ^18^). Surprisingly, RASSF1C expression is maintained in tumors and emerging studies indicate a pro-oncogenic role for this isoform of the *RASSF1* gene ^22^. We have previously linked RASSF1C to SRC kinase activation and aggressiveness in breast cancer, contributing to explain the biological events leading to the aggressive phenotype observed in *RASSF1* A-methylated tumors ^17^. To study the effect of RASSF1C oncoprotein (Supp. Fig. 1A) in breast cancer we employed MCF7 and MDA-MB-231 cell lines, both lacking expression of the tumor suppressor isoform RASSF1A, as shown in reports from our and other groups ^17,23–25^. We first assessed RASSF1C transcript endogenous levels in both breast and H1299 lung cancer cell lines (Supp. Fig. 1B, left). The mRNA levels of RASSF1C were significantly higher in MDA-MB-231 cells compared to RASSF1A transcript levels (Supp. Fig. 1B, middle), in agreement with reports indicating loss of RASSF1A expression in this cell line. Additionally, we confirmed expression levels of RASSF1C mRNA after MDA-MB-231 cells were transfected with a plasmid encoding for a control sequence (pcDNA3) or RASSF1C (Supp. Fig. 1B, right graph).

The process of mesenchymal-amoeboid transition (MAT) is associated with changes in cell morphology, where actin cytoskeleton and stress fibers are reorganized into a contractile actomyosin cortex. Cancer cells adopting amoeboid mode of motility have then typical rounded morphology in three-dimensional matrices. To determine whether RASSF1C is involved in MAT, we transiently transfected MCF7 cells with a fluorescently tagged ZsGreen-RASSF1C (ZsRASSF1C) or control construct (ZsGreen). We observed that ZsRASSF1C expressing cells grown on 2D surface displayed a progressive decrease of diameter within 12 h of expression of the construct compared to control (Fig. 1a, Supp. Fig. 1C, Video 1). As rounded phenotype is a morphological change associated with reorganization of actin cytoskeleton and actin cortical localization, this indicated a potential correlation between RASSF1C expression and mesenchymal-amoeboid transition (Fig. 1a). In order to verify whether these phenotypic changes could also be observed in three-dimensional matrices, the necessary 3D environment for upregulation of Rho/ROCK signaling, we cultured RASSF1C expressing cells in 3D extracellular collagen matrices. We observed increased numbers of RASSF1C expressing cells that exhibited rounded morphology in 3D collagen matrix in mesenchymal (MDA-MB-231) (Fig. 1b) and also in epithelial (MCF7 and H1299) cell lines (Fig. 1b, Supp. Fig. 1D, E), suggesting that RASSF1C may be involved not only in mesenchymal-amoeboid but also in epithelial-amoeboid transition. Next we visualized the co-localization of f-actin (via phalloidin stain) and pMLCII, a marker of high myosin II activity correlated to amoeboid morphology ^26^, which indicated phenotypic shift from actin stress fibers in the mesenchymal control cells (visible in the phalloidin stain, Fig. 1c top panel), to cortical actomyosin in RASSF1C-expressing MDA-MB-231 cells (visible in both pMLCII and phalloidin stain, Fig. 1c bottom panel). Therefore, as RASSF1C-mediated morphological changes are associated with cortically localized actin and myosin, this indicated MAT ^27,28^. To additionally confirm whether RASSF1C cells also actively adopt rounded, amoeboid phenotype during MAT process, we produced MDA-MB-231 spheroids and embedded them into a 3D collagen matrix. We observed that single cells in both control (ZsGreen) and ZsRASSF1C^+ve^ 3D spheroids were able to actively detach and migrate from the spheroids. However, while control cells maintained a typically elongated, fibroblast-like, shape, cells transfected with RASSF1C showed rounded morphology (Supp. Fig. 1F). We reasoned that expression of RASSF1C is responsible for epithelial-amoeboid (EAT, in MCF7 and H1299 cell lines) and mesenchymal-amoeboid transition (MAT, in MDA-MB-231 cell line), and for the switch from elongated into rounded morphology induced in 3D collagen matrix. To corroborate these results, we tested whether loss of RASSF1C can revert the phenotype from amoeboid to mesenchymal. To this end, suppression of endogenous RASSF1C expression via siRNA mediated knockdown in MDA-MB-231 cells resulted in a significant decrease in the baseline number of amoeboid cells (displaying rounded morphology) and a significant increase in the number of mesenchymal cells (displaying elongated phenotype) in 3D matrix (Fig. 1d).

**Figure 1.**
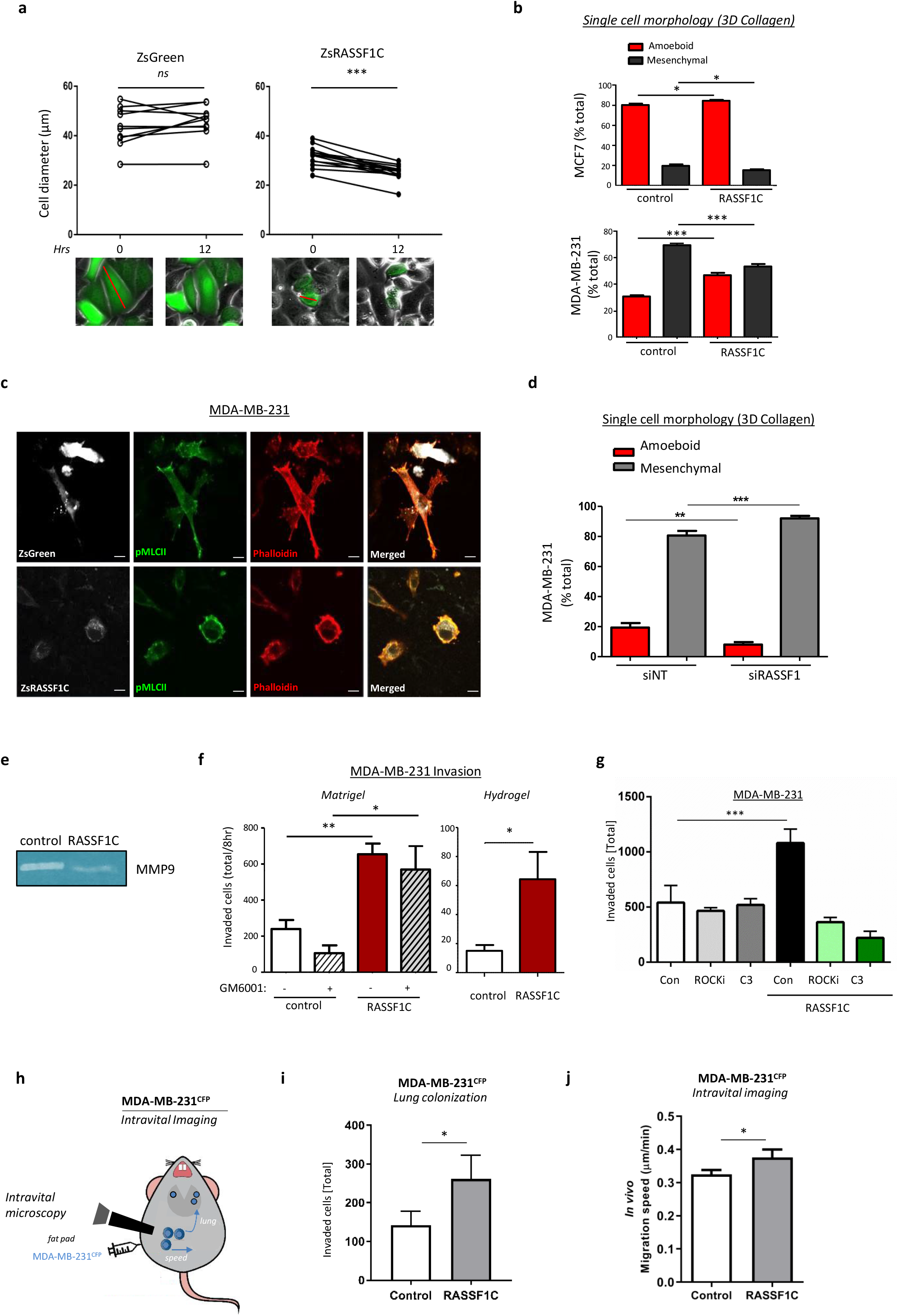
a. Top: graphs showing the change in cell diameter for MCF7 cells transfected with ZsGreen (left) or ZsRASSF1C (right) plasmids at 0 h and 12 h, connections indicate individual cell tracking. Bottom: example images diameter measurements (red line). b. Single cell morphology of MCF7 and MDA-MB-231 cell lines, expressing or not a RASSF1C construct, cultured in 3D rat tail collagen I and analyzed for their amoeboid (rounded) or mesenchymal (fibroblast-like) morphology 24 hours after seeding in 3D matrix. c. Confocal images of MDA-MB-231 cells transfected with ZsGreen or ZsRASSF1C grown in 3D-collagen matrix. Cells were imaged and stained with Phalloidin-568 (red), pMLCII/Alexa-633 (false-coloured green) or imaged for ZsGreen (false-coloured white). Scale 20 μm. d. Single cell morphology assay of MDA-MB-231 cells cultured in 3D-collagen and analysed for morphology 24 hours after transient knock-down using either a control sequence (NT, non-targeting) or a sequence targeting the *RASSF1* locus. RASSF1 knock-down marks a reduction in amoeboid cells. e. Gelatine zymography assay shows downregulation of MMP-9 metalloprotease in MDA-MB-231 cells over/expressing RASSF1C. f. 3D Matrigel Boyden chamber invasion with or without metalloproteases inhibitor GM6001 and 3D Hydrogel invasion of MDA-MB-231 control cells or cells over-expressing RASSF1C, (24 h). g. Quantification of Boyden chamber invasion after 24 hours in 3D matrigel of MDA-MB-231 cells, transfected with ZsGreen or ZsRASSF1C and treated with inhibitors against ROCK (Y-27632, 10 μmol/L) or Rho (C3, 2 μg/ml). h. Cartoon summarising intravital imaging experiments. i. Lung colonization of MDA-MB-231 CFP^+^ cells (MDA-MB-231^CFP^) stably expressing control or HA-RASSF1C in mammary gland tumors from 5 mice each, measured from intravital imaging time-lapse images. j. Average migration speed (expressed as μm/min) of n=30 randomly selected CFP positive cells in intravital images of Control and RASSF1C expressing MDA-MB-231 tumors. All data are from n=3 independent experiments. Data are represented as mean ± SEM.* p ≤ 0.05, ** p ≤ 0.01, *** p ≤ 0.001.

As reported in the literature, mesenchymal invasion is dependent on production of metalloproteases that mediate proteolytic degradation of the extracellular matrix ^29^. Conversely, amoeboid invasion does not rely on proteolytic degradation of surrounding tissue ^30^, but is dependent on either squeezing through pre-existing pores in the extracellular matrix or deforming surrounding tissue by tension generated by cortical actomyosin ^28,31^. To investigate the production of metalloproteases we performed gelatin zymography assay *in situ*, where we analyzed MMP enzymatic activity from the cell medium ^32^. We detected increased activity of MMP9 metalloprotease in control MDA-MB-231 cells compared to cells expressing RASSF1C (Fig. 1e). To confirm this observation, we performed Boyden chamber invasion assay with a broad spectrum metalloprotease inhibitor, GM6001. As expected, in control cells GM6001 inhibited matrigel invasion (Fig. 1f, left bar graph). However, in the presence of RASSF1C, metalloprotease inhibition did not influence the ability of cells to invade matrigel (Fig. 1f, left bar graph). These data suggested that RASSF1C-mediated mode of invasion was not dependent on metalloproteases. To further investigate this, we performed an invasion assay using hydrogel matrix, which is not degradable by metalloproteases (Fig. 1f, right bar graph, and Supp. Fig. 1G). Hydrogel invasion assay showed that control mesenchymally invading cells were not able to invade the hydrogel matrix, whereas RASSF1C cells ability to invade the matrix was significantly greater (Fig. 1f, right bar graph). These results strongly support the hypothesis that RASSF1C-mediated mode of invasion is metalloprotease-independent. Thus, we reasoned that invasion may be supported by traction forces generated via contractile actomyosin in a three-dimensional environment. Given that Rho/ROCK signaling is the major pathway involved in amoeboid motility ^33^, we next asked whether the observed RASSF1C-promoted amoeboid invasion is a consequence of Rho/ROCK upregulation in 3D. Using ROCK (Y-27632) and Rho (C3) inhibitors we could demonstrate that matrigel invasion by ZsRASSF1C-expressing MDA-MB-231 cells was greatly impaired by both inhibitors, thus demonstrating Rho/ROCK dependency (Fig. 1g and Supp. Fig. 1H). However, the treatment of control cells with both drugs resulted in no effect on their ability to invade (Fig. 1g and Supp. Fig. 1H). These data confirmed our previous results, showing that MDA-MB-231 control cells use a mesenchymal mode of invasion which is dependent on metalloprotease degradation of ECM (Fig. 1f). To further support Rho/ROCK dependency, we could show that expression of ZsRASSF1C and a dominant negative RhoA derivative (RhoA-DN) suppressed ZsRASSF1C-driven invasion in both MDA-MB-231 and MCF7, while a catalytically active RhoA derivative (RhoA-CA) enhanced invasion upon RASSF1C expression (Supp. Fig. 1I, J).

We next set out to determine whether RASSF1C could promote amoeboid invasion and metastatic spread *in vivo.* To this end, we employed CFP-labelled MDA-MB-231 cells (MDA-MB-231^CFP^, ^34^) and stably expressed RASSF1C to promote a switch to amoeboid motility (MDA-MB-231^CFP;HA-RASSF1C^). Tumors were initiated in the mammary gland of immuno-compromised mice, and lung colonization as well as migration of individual cells was tracked by time-lapse images via intravital microscopy (IVM) (Fig. 1h). Importantly, we observed that the number of MDA-MB-231^CFP;HA-RASSF1C^ cells that colonized distal sites in the lungs was significantly greater than the number of control cells, therefore supporting RASSF1C oncogenic potential *in vivo* (Fig. 1i). Low-adhesion attachment during *in vitro* amoeboid invasion in 3D matrices allows amoeboid cells to translocate at relatively high velocities, between 2 to 25 μm/min, while relative speed of mesenchymally invaded cells is approximately 0.1-0.5 μm/min due to recruitment of proteolytic enzymes and slow turnover of focal adhesions ^33,35,36^. We next investigated the speed adopted by RASSF1C cells during *in vivo* invasion. Manual analysis of the invading cell speed in MDA-MB-231^CFP;HA-RASSF1C^ and MDA-MB-231^CFP,Control^ tumors showed higher migration speed of cells in MDA-MB-231^CFP;HA-RASSF1C^ tumors (Fig. 1j). Interestingly, these cells adopted more amoeboid, rounded morphology during their invasion from MDA-MB-231^CFP;HA-RASSF1C^, compared to control tumors (Supp. Fig. 1K, morphology was defined as amoeboid, hybrid or mesenchymal).

The data so far suggest that RASSF1C expression promotes a phenotypic switch from mesenchymal or epithelial to amoeboid morphology and motility both *in vitro* and *in vivo*, likely through upregulation of Rho/ROCK/pMLCII pathway.

### RASSF1C-mediated SRC activation promotes pro-amoeboid Rho/ROCKI/pMLCII signaling

We next set out to explore the mechanism through which RASSF1C promotes mesenchymal-amoeboid transition (MAT). RASSF1C directly supports activation of SRC kinase by preventing CSK inhibitory phosphorylation on Y527 ^17^. Notably, SRC activation is well documented to promote mesenchymal motility, however, it can also facilitate contractility via inhibitory phosphorylation of RhoGDI (pY156), which results in activation of RhoA ^21^. Therefore, we hypothesized that RASSF1C activation of SRC may be implicated in MAT via upregulation of Rho/ROCK/pMLCII signaling pathway. As the Rho interacting domain of RASSF1A is identical to RASSF1C ^14^, we investigated whether RASSF1C promotes Rho activation. Rho-GTP pull-down assay on lysates obtained from cells expressing a control or a RASSF1C construct showed that RASSF1C expression results in accumulation of active Rho (shown as Rho-GTP, MDA and MCF7 cells (Fig. 2a). Next, we performed Rho-GTP pull down in cells expressing RASSF1C and co-expressing a SRC construct or depleted of SRC by siRNA knock-down (Fig. 2b). Transient knock-down of SRC by siRNA showed a decrease in Rho activity (measured as lower expression levels of Rho-GTP in the pull-down). Interestingly, SRC over-expression resulted in increased inhibitory phosphorylation of RhoGDI (pY156), a Rho GTPase inhibitor, and elevated levels of pMLCII (pS19) (Fig. 2b). Similarly, RASSF1C over-expression alone could induce RhoGDI phosphorylation, increased expression of ROCKI and pMLCII (Supp. Fig. 2A). Furthermore, inhibition of endogenous RASSF1C expression by siRNA in MDA-MB-231 cells resulted in reduced SRC activating phosphorylation (pY416), increased SRC inhibitory phosphorylation (pY527), concomitant with increased expression of β1-integrin, a crucial regulator of stable attachment to ECM in mesenchymally invaded cells ^30^ (Fig. 2c). The above results led us to hypothesize that the morphological changes observed in RASSF1C cells (Fig. 1) were due to RASSF1C upregulation of Rho/ROCK/pMLCII signalling axis via activation of SRC. SRC in turn inhibits RhoGDI, event that allows activation of Rho. To confirm that SRC and RhoGDI are downstream of RASSF1C in Rho/ROCK upregulation, we performed co-immunoprecipitation and determined that both RASSF1C and RhoGDI are found in complex with SRC (Fig. 2d, 2e cartoon that recapitulates the hypothesized mechanism). A previous study described that the main isoform of the *RASSF1* gene, the tumor suppressor RASSF1A, is able to repress RhoA activity and MLCII phosphorylation through its RA (Ras association) domain ^14^. Based on this report, we next generated two point mutations in the identical RA domain sequence of RASSF1C, RASSF1C-R197W and RASSF1C-R199F, which had previously been demonstrated to be necessary for RASSF1A-mediated inhibition of RhoA ^14^. Compared to RASSF1C, RASSF1C-R199F and RASSF1C-R197W mutants were less capable of stimulating SRC binding to RhoGDI or activate Rho, which in turn led to decreased MLCII phosphorylation and ROCKI expression (Fig. 2f, g and Supp. Fig. 2B). Moreover, we confirmed that RASSF1C-R199F and RASSF1C-R197W mutants failed to promote rounded morphology, and instead exhibited elongated, fibroblast-like morphology in 3D collagen matrix, typical of mesenchymal cells (Fig. 2h, i and Supp. Fig. 2C). This mesenchymal phenotype was accompanied by formation of abundant stress fibers with no signs of formation of cortical actomyosin (Fig. 2i). Tension and contractility of the actomyosin network initiated by Rho/ROCK signaling in amoeboid cells promote rupture of the actomyosin cortex resulting membrane blebs, spherical membrane herniations ^27^. These membrane blebs were typical only in RASSF1C amoeboid cells, while RASSF1C-R199F and RASSF1C-R197W mutants maintained their mesenchymal, elongated phenotype in 3D substrates (Supp. Fig. 2D). Here we show for the first time, that pro-mesenchymal activated SRC can also be involved in amoeboid invasiveness via inhibition of RhoGDI, a RhoA antagonist, which in turn allows activation of Rho/ROCK/pMLCII signaling and cytoskeleton reorganization.

**Figure 2.**
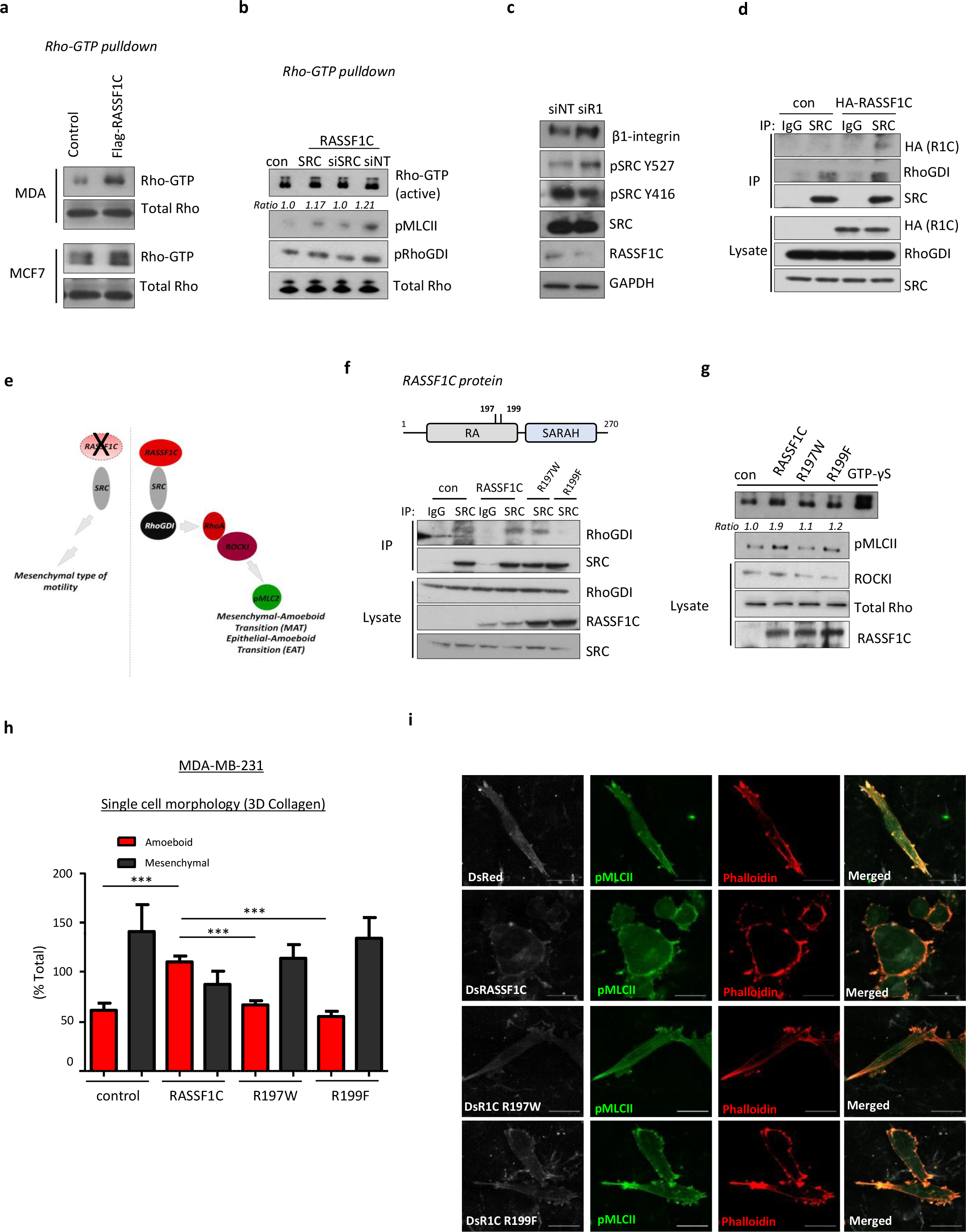
a. Western blot analysis of Rho-GTP pull-down assay in MDA-MB-231 and MCF7 cell lines transiently transfected either with control or Flag-RASSF1C plasmid. b. Rho-GTP pull-down binding assay in MDA-MB-231 control or RASSF1C over-expressing cells and cotransfected either with SRC plasmids, siRNA against SRC or siNT (non-targeting sequence). Western blot analysis shows Rho activity and pMCLII and pRhoGDI binding. c. Western blot analysis of proteins in MDA-MB-231 cells transiently transfected with siRNA targeting a control sequence (NT) or targeting RASSF1. GAPDH was used as a loading control. d. Immunoprecipitation from MDA-MB-231 transiently transfected with control (pcDNA3) or HA-RASSF1C plasmid, pulled down using a SRC antibody or same species IgG antibody. e. Schematic cartoon recapitulating the proposed RASSF1C/SRC-driven mechanism. f. Top, schematic of RASSF1C domain structure indicating putative mutations in the RA domain affecting RhoA activity. Bottom, immunoprecipitation of MDA-MB-231 transiently transfected with either control (DsRed), DsRASSF1C plasmid, DsRASSF1C-R197W or DsRASSF1C-R199F mutants. Pull-down was performed using a SRC or same species IgG antibody and blotted with indicated antibodies. g. Western blot analysis of Rho-GTP pull-down assay in MDA-MB-231 cells transiently transfected with RASSF1C or its mutants (R197W, R199F). h. Quantification of amoeboid versus mesenchymal cells in single cell morphology assay in 3D-collagen indicating the degree of mesenchymal-amoeboid transition of MDA-MB-231 cells when RASSF1C or the described mutants are expressed. i. Representative confocal images show mesenchymal or amoeboid cells per field of view of MDA-MB-231 cells transfected with DsRed, DsRASSF1C, DsRASSF1C-R197W or DsRASSF1C-R199F, grown in 3D-collagen and stained with Phalloidin-568 (red) and pMLCII/Alexa 633. Scale bars represent 10 μm. For both Rho-GTP pull-downs and immunoprecipitation assays total proteins were used as loading controls. All data are from n=3 independent experiments. Data are represented as mean ± SEM.* p ≤ 0.05, ** p ≤ 0.01, *** p ≤ 0.001.

### RASSF1C amoeboid cells release pro-invasive Extracellular Vesicles

Having shown that RASSF1C expression leads to mesenchymal to amoeboid transition (MAT) and invasion via generation of protrusive forces by upregulation of the Rho-ROCK signaling pathway, we hypothesized that RASSF1C could promote invasion via additional mechanisms. Growing evidence in the literature describes the importance of extracellular cues, such as extracellular vesicles (EVs), involved in intercellular communication and in mediating cancer invasiveness ^37^. To study cell-cell communication via EV transfer we employed a previously reported system comprised of a donor and a reporter cell line. The donor cell line, i.e. MDA-MB-231^CFP,UbC-Cre^, expressing Cre under the ubiquitin promoter, has been shown to package Cre mRNA into EVs ^34^. The reporter cell line, i.e. T47D luminal breast cancer cells, harbors both a floxed *DsRed* and a lox-stop-lox *eGFP* allele. When the two cell types are co-cultured, T47D reporter cells that internalize EVs containing Cre mRNA switch from DsRed to eGFP, thereby allowing distinction between Cre^+^ EV recipients (green) from unrecombined T47D cells (red, Fig. 3a, left). We tested the system *in vitro* by generating donor MDA-MB-231^CFP,UbC-Cre^ cells stably expressing a control or a RASSF1C construct (Supp. Fig. 3A). We performed experiments with reporter T47D cells where donor/reporter cells were either in direct contact (co-culture) or separated by a 0.4 μm porous membrane (transwell). Both the co-culture (Supp. Fig. 3B) and the transwell setup (Fig. 3a) showed a significant increase in eGFP events, thereby indicating that RASSF1C modulates cell communication, likely through transfer of EVs.

**Figure 3.**
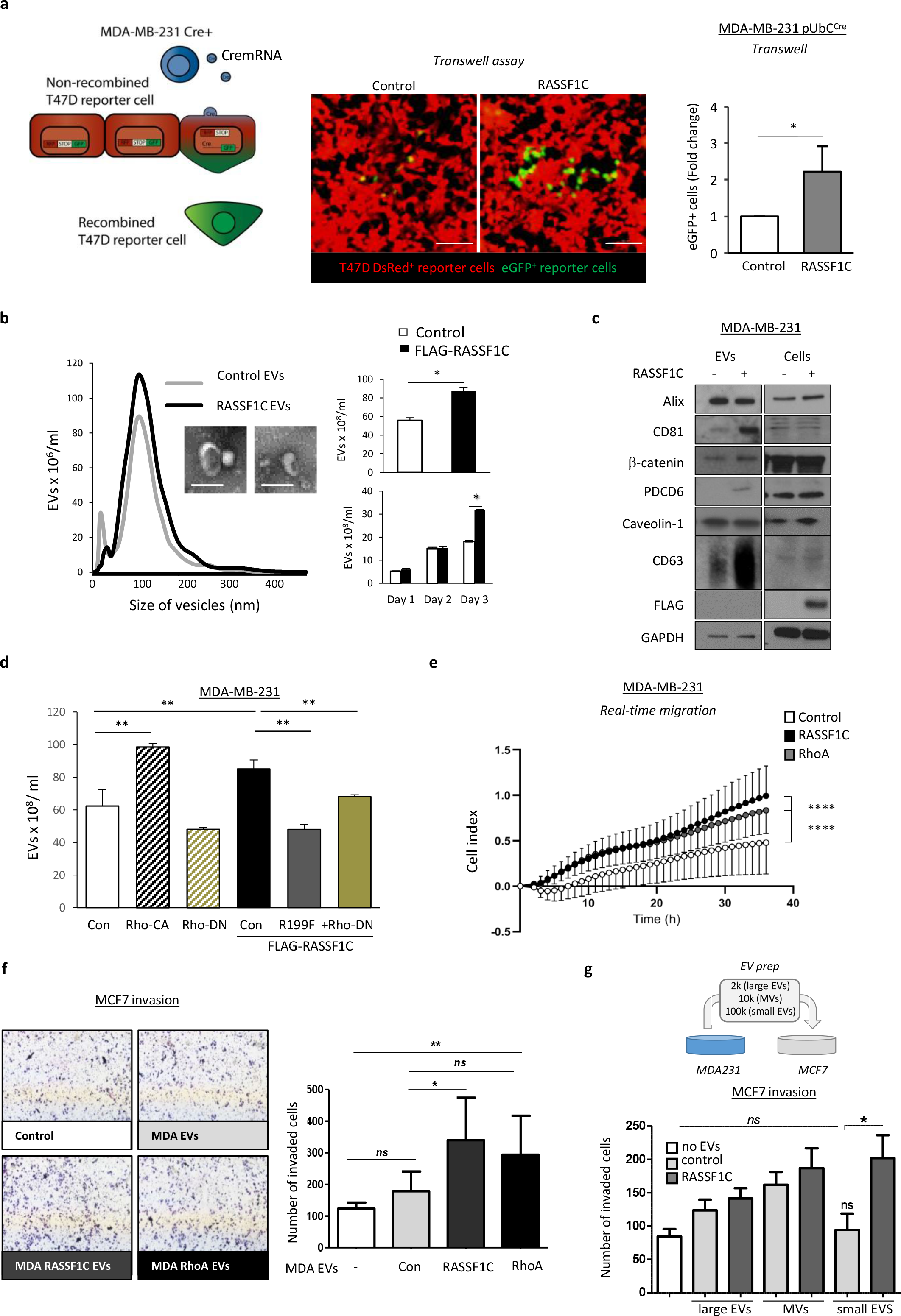
a. Left: Cartoon showing the principle behind the Cre-carrying EVs and how recipient cells switch from DsRed^+^ to eGFP^+^ expression upon Cre-mediated excision of the DsRed locus. Centre: representative images, and right: quantification of T47D^DsRed^ cells that converted to eGFP^+^ in the bottom wells of a transwell system in the presence of MDA-MB-231^CFP;Cre;Control^ or MDA-MB-231^CFP;Cre;HA-RASSF1C^ seeded in the upper wells. b. Left: averaged Nanoparticle Tracking Analysis displaying the size distribution of EVs isolated from media of MDA-MB-231 cells expressing Control (pcDNA3) or FLAG-RASSF1C plasmid. Inserts: transmission electron microscopy images of single EVs. Scale 200 nm. Right: quantification of total secreted EVs (96 h, top) or at indicated times (below), as measured by NTA. c. Western blot analysis of EVs isolated from MDA-MB-231 cells transfected with Control (pcDNA3) or FLAG-RASSF1C with indicated antibodies. Equal amounts of protein were loaded after normalization via microBCA assay. d. Quantification of EVs isolated from MDA-MB-231 cells transfected with Control (pcDNA3), RhoA-CA, RhoA-DN, FLAG-RASSF1C, FLAG-RASSF1C-R199F, or FLAG-RASSF1C + RhoA-DN using Nanoparticle Tracking Analysis. e. Normalized values of real-time invasion rates in xCELLigence plates of MDA-MB-231 cells treated with 20 ng/μl EVs derived from MCF7 cells expressing a Control, FLAG-RASSF1C, or EGFP-RhoA plasmid, left to migrate for 36 h. f. Left: Representative images of Boyden chamber invasion assay in 3D matrigel of MCF7 cells, treated with 20 ng/μl EVs derived from MDA-MB-231 cells, transfected the same constructs used in (e). Right: Quantification of Boyden chamber invasion 24 hours after EV treatment. g. Top: graphical representation of the functional (invasion) assay performed using MDA-MB-231 derived EVs on MCF7 recipient cells. Bottom: Boyden chamber invasion assay in 3D matrigel of EVs isolated from different steps of ultracentrifugation. EVs were collected from conditional medium produced by MDA-MB-231 expressing Control (pcDNA3) or FLAG-RASSF1C. The 2K (large EVs), 10K (medium EVs or MVs) and 100K (small EVs, including exosomes) pellets were collected and equal amount of EVs (normalized according to protein content) were incubated with MCF7 cells. MCF7 were evaluated for invasion after 24 hours. All data are from n=3 independent experiments. Data are represented as mean ± SEM.* p ≤ 0.05, ** p ≤ 0.01, *** p ≤ 0.001, **** p ≤ 0.0001.

In line with these results, we therefore sought to understand whether amoeboid RASSF1C cells shed EVs able to transfer the invasive potential to recipient cells. We isolated EVs, via a differential centrifugation protocol, from MDA-MB-231 cells expressing either a control or Flag-RASSF1C construct, cultured in 0.1% EV-depleted FBS. Employing Nanoparticle Tracking Analysis (NTA) we observed a significantly higher number of EVs released by RASSF1C cells (Fig. 3b), which could explain the differences observed in recombined reporter cell numbers in the Cre/LoxP model (Fig. 3a). The mean EV size (~ 100 nm) indicated isolation of small EVs and it was confirmed by Electron Microscopy (Fig. 3b insets and Supp. Fig. 3C for complete field of view). In order to rule out the possibility that exogenous plasmid expression alone could drive EV release, we cultured cells for up to 72 h and isolated EVs at different time-points (24, 48 and 72 h, Fig. 3b, bottom bar graph). Collection of the conditioned media at defined time-points suggested that the accumulation of EVs occurred over days following transfection with RASSF1C, suggesting that this occurs in response to changes in cell behavior and is not an acute release in response to RASSF1C expression. We next set out to characterize released EVs by protein composition. Western-blotting of isolated EVs showed selective enrichment of some commonly used EV markers, e.g. CD63 and CD81 vs Alix, in RASSF1C samples (Fig. 3c). Interestingly, RASSF1C protein enrichment on EVs was not observed (Fig. 3c, Flag tag). In order to obtain a more comprehensive picture of the proteome present in RASSF1C derived EVs we performed Mass Spectrometry and found that the general proteome of MDA-MB-231 EVs is dominated by proteins involved in integrin signaling and cytoskeletal regulation by Rho signaling (PANTHER pathway enrichment, Supp. Fig. 3D), but not greatly altered by RASSF1C. This observation suggests that any functional effect of RASSF1C-derived EVs may be ascribed to a different type of EV cargo or to a specific EV subpopulation. Having shown that RASSF1C expression leads to amoeboid transition in MDA-MB-231 cells, we next sought to understand if Rho signaling was also involved in RASSF1C-mediated release of EVs. Previously used RhoA and RASSF1C constructs (Supp. Fig. 1I, Fig. 2f-h) were expressed in MDA-MB-231 cells and EVs were isolated as described before. NTA measurements showed that over-expression of a constitutively active derivative of RhoA (RhoA-CA) enhanced EV release similarly to RASSF1C expression, whereas a RhoA dominant negative derivative (RhoA-DN) had an opposite effect (Fig. 3d). Moreover, expression of a RASSF1C derivative with impaired capacity of activating Rho signaling (RASSF1C-R199F) or co-expression of RASSF1C wild-type and RhoA-DN also hinders EV release, suggesting that the effects on EVs driven by RASSF1C may involve RhoA activation.

Having shown that RASSF1C cells promote EV transfer, we next asked whether EVs can confer a selective migratory and invasive advantage to recipient cells. In order to answer this question, we first performed an uptake experiment to ensure that isolated EVs are functional and can be further used for functional assays. EVs were isolated from MDA-MB-231 as previously described, labelled with PKH67 lipophylic dye and equal amounts of EVs (normalized for protein content by microBCA) were incubated with recipient MDA-MB-231 cells. As shown in Supp. Fig. 3E, both control and RASSF1C EVs were internalized by recipient cells at similar rates.

We next performed functional assays by incubating recipient (MDA-MB-231 or MCF7) cells with EVs isolated from donor MDA-MB-231 cells expressing a control, a RASSF1C or a RhoA construct.

Incubation of cells with RASSF1C or RhoA EVs (20 ng/μl dose) stimulated significantly higher cell migration than incubation with control EVs (Fig. 3e, displayed results are normalized to baseline untreated cells). At higher doses (40 ng/μl, Supp. Fig. 3F), EVs derived from control cells were able to promote recipient cell migration at similar rates to EVs derived from RASSF1C or RhoA.

We next incubated the same panel of EVs with recipient MCF7 cells seeded on a matrigel layer, in order to measure matrigel invasion. Strikingly, the number of invaded cells was significantly higher when cells were treated with 20 ng/μl RASSF1C or RhoA EVs, compared to control EVs and untreated cells (Fig. 3f, Supp. Fig. 3G for a comparison of both concentrations).

Cells have been shown to release heterogeneous populations of EVs ^38^ and classification of the different subgroups is still subject to ongoing research ^39^. In order to investigate whether multiple EV populations could transfer the observed oncogenic phenotype to recipient cells, we isolated fractions using three centrifugation steps: 2,000×*g* (large EVs), 10,000x*g* (MV or medium EVs, ^12^), and 100,000x*g* (small EVs, among which exosomes exist, corresponding to the protocol used throughout the study), and incubated recipient cells with control or RASSF1C EV populations (Fig. 3g). Fig. 3g shows that only RASSF1C small EVs induced a significantly higher number of cells to invade matrigel in a transwell assay, if compared to control small EVs. In our setup, the RASSF1C large and medium EVs did not confer higher invasive ability to recipient cells, compared to the corresponding control fractions.

To further establish that the small EVs population is mediating the invasive phenotype observed in the small EV fraction recovered from ultracentrifugation (Fig. 3g), we employed a Tangential Flow Filtration/Size Exclusion Chromatography isolation protocol and incubated recipient cells with RASSF1C or control small EVs (Supp. Fig. 3H). First, EVs isolated with this method were characterized by western-blotting and NTA. EVs displayed protein markers already detected with the ultracentrifugation protocol (Supp. Fig. 3H, left) and NTA showed similar particle number between RASSF1C and control EVs, with this isolation method (Supp. Fig. 3H, middle). Importantly, RASSF1C small EVs still induced a significantly higher number of recipient cells to invade the matrigel membrane, compared to control small EVs (Supp. Fig. 3H, right). The data so far suggest that RASSF1C-derived small EVs, and no other fractions, transfer the invasive potential to naïve cells.

It is important to note that, throughout our study, the treatment of recipient cells with MDA-MB-231 derived control EVs (expressing a control vector) already induce an increase in cell migration or invasion, compared to untreated cells (Fig. 3e-g, Supp. Fig. 3H). This is expected, as it is well-documented that EVs from highly invasive cancer cells can promote migration and invasion in recipient cells. However, oncogenic activity of RASSF1C consistently led to a significant increase of invasive potential in recipient cells.

Taken together, the data show that RASSF1C promotes EV functional transfer and that both RASSF1C and RhoA expression promote the release of pro-migratory and pro-invasive EVs. Our data suggest that RASSF1C oncogenic potential has an effect on EV secretion and this may occur through Rho activation.

### RASSF1C amoeboid cells display cancer stemness features transferrable via EVs

Amoeboid transition has been previously linked to cancer stemness in melanoma cells and RhoA has been directly associated to the expression of stem-like features ^40,41^. Given the oncogenic role of RASSF1C and its ability to activate Rho (Fig. 2), we next assessed whether RASSF1C-promoted mesenchymal-amoeboid transition could also be associated with cancer stemness during *in vitro* invasion. A plethora of markers have been elucidated and used to validate a cancer stemness profile. Among those, the surface glycoprotein CD133 and the aldehyde dehydrogenase enzyme ALDH1 are currently used as breast cancer stemness markers ^42^. We therefore assessed the expression of ALDH1 and CD133 in RASSF1C three-dimensional spheroids cultivated on matrigel matrix to mimic colony formation assay, an assay widely used to investigate stemness phenotype *in vitro.* We observed significantly higher expression of these markers in RASSF1C spheroid colonies, compared to control 3D spheroids (Fig. 4a). NANOG, SOX2 and OCT4 (octamer-binding transcription factor 4) are key regulators of pluripotency and tumor invasion and they have been proposed as biomarkers for cancer stemness in breast and renal carcinoma ^43,44^. MDA-MB-231 spheroids expressing either a ZsGreen or ZsGreen-RASSF1C construct were embedded in 3D collagen matrix, cells were allowed to invade from the aggregates and stained for the stemness marker Nanog. A Z-stack image in Fig. 4b shows that invaded amoeboid cells from ZsGreen-RASSF1C spheroids retained both rounded morphology (see magnification in Fig. 4b) and also higher Nanog expression compared to control. However, migrating cells from control 3D spheroids exhibit reduced Nanog fluorescence intensity, suggesting that mesenchymal invasion is not associated with a stemness phenotype. Furthermore, transient knock-down of RASSF1C decreased CD133 protein levels (Fig. 4c), as well as fluorescent intensity of ALDH1 and Nanog in MDA-MB-231 cells (Fig. 4d and Supp. Fig. 4A).

**Figure 4.**
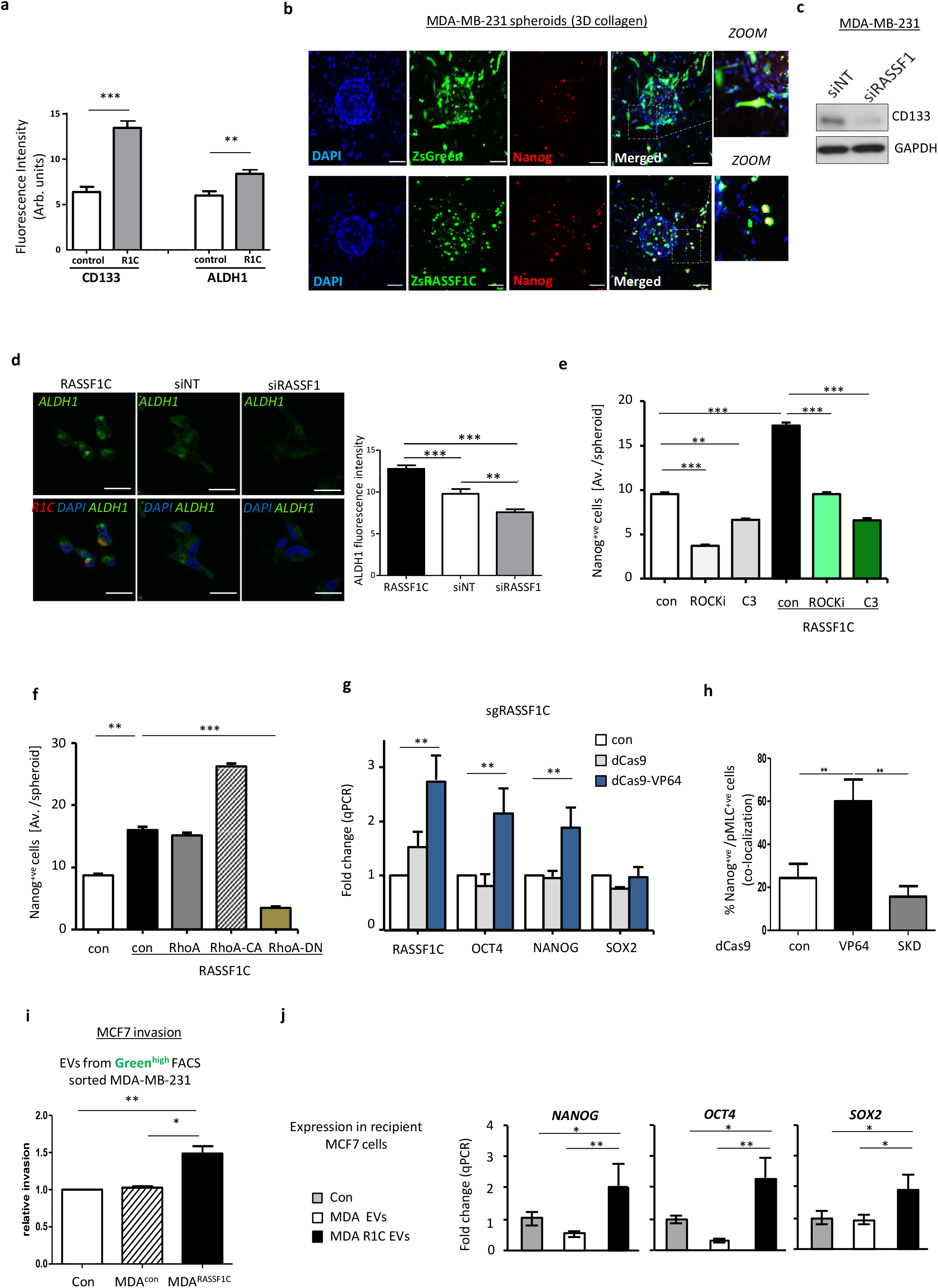
a. Quantification of fluorescent intensity of CD133 and ALDH1 cancer stem cell markers in MDA-MB-231 control or RASSF1C transfected spheroids cultured on Matrigel matrix. b. Confocal images of ZsGreen or ZsRASSF1C expressing MDA-MB-231 spheroids embedded in 3D rat tail collagen I. Cells were stained with Nanog (red). Zoom images show that RASSF1C expressing cells are also Nanog positive. Scale 100 μm. c. Western blot analysis of CD133 protein expression level in MDA-MB-231 cells treated with siNT or siRASSF1. d. Representative confocal images (upper row) and quantification of fluorescent intensity of ALDH1 expression in MDA-MB-231 cells expressing RASSF1C (black bar in the graph) or after siRNA treatment against RASSF1 (grey bar). Lower row shows merged images with DAPI (blue), ALDH1 (green) and RASSF1C (red). Scale bars represent 20 μm. e. Quantification of the absolute number of Nanog positive cells (measured as average cell number/spheroid) per MDA-MB-231 ZsGreen or ZsRASSF1C spheroids in 3D-collagen, treated with ROCK inhibitor (Y-27632, 10 μmol/L) or Rho inhibitor (C3, 2 μg/ml). f. Quantification of the absolute number of Nanog positive cells (measured as average cell number/spheroid) per MDA-MB-231 ZsGreen or ZsRASSF1C spheroids in 3D-collagen, co-transfected with either RhoA WT, catalytically active (CA) or dominant negative (DN) constructs. g. Fold change in qRT-PCR of Nanog, OCT4 and SOX2 mRNA from MCF7 cells expressing CRISPR-dCas9 or dCas9-VP64 plus single guide RNA for the RASSF1C promoter (sgRASSF1C). h. Quantification of the percentage of double positive pMLCII and Nanog cells in MCF7 cells expressing CRISPR-dCas9, dCas9-VP64, dCas9-SKD plus single guide RNA for the RASSF1C promoter (sgRASSF1C). At least 300 cells were counted per experiment. i. Quantification of number of invaded MCF7 cells in 3D matrigel, after incubation with EVs from MDA-MB-231 cells sorted for Green^high^ expression. MDA-MB-231 cells stably expressing a pQXCIP-zsGreen or pQXCIP-zsGreen-RASSF1C construct were sorted and EVs were harvested 24 h later, prior to seeding with MCF7 cells in Boyden chambers. Invasive potential of MCF7 cells was evaluated 24 h after incubation with EVs. j. qRT-PCR expression of NANOG, OCT4, SOX2 in MCF7 cells 24 h after exposure to 50 ng/μl EVs derived from MDA-MB-231 cells (treated as indicated). All data are from n=3 independent experiments. Data are represented as mean ± SEM.* p ≤ 0.05, ** p ≤ 0.01, *** p ≤ 0.001.

To verify that cancer stemness is dependent on amoeboid Rho/ROCK signaling, we treated control and RASSF1C spheroids in 3D collagen matrix with either a RhoA (C3) or a ROCK inhibitor. Both inhibitors significantly decreased the number of Nanog^+^ cells in ZsGreen-RASSF1C expressing spheroids (Fig. 4e and Supp. Fig. 4B, C). Similarly, co-expression of RASSF1C and RhoA-CA constructs greatly increased Nanog expression in MDA-MB-231 3D spheroids, however, this effect was reverted when RhoA-DN was co-expressed (Fig. 4f), suggesting Rho/ROCK dependency. Furthermore, RASSF1C-R199F mutant cells, cultured in 3D collagen matrix also showed impaired Nanog expression, due to inability of Rho activation by this mutant (Supp. Fig. 4D). In order to ensure that the expression of stem-like features was originating from the RASSF1C-expressing pool of cells, we sorted for green^high^ expression MDA-MB-231 stably expressing ZsGreen or ZsGreen-RASSF1C and analyzed mRNA levels of pluripotency genes such as NANOG, OCT4 and SOX2. All three markers were significantly upregulated in sorted ZsGreen-RASSF1C cells (Supp. Fig. 4E). In order to validate RASSF1C as a modulator of cancer stemness with an alternative approach, we induced endogenous *RASSF1C* expression by epigenetic editing; where inactive Cas9 derivatives (dCas9) fused to a transcriptional activation domain (dCas9-VP64) and single guide RNAs target the internal *RASSF1* promoter (sgRASSF1C) to selectively promote expression of the RASSF1C transcript (see Supp. Fig. 4F, graphical abstract). MCF7 cells expressing a dCas9-VP64 construct showed upregulation of NANOG and OCT4, along with the RASSF1C mRNA transcript (Fig. 4g). Interestingly, in this assay, no upregulation of the pluripotency marker SOX2 was observed. Next, we aimed to show that RASSF1C amoeboid cells and cancer stem cells are the same entities. To this end, we employed the Cas9 technique and visualized both pMLCII (an amoeboid cell marker) and Nanog (a cancer stem cell marker). Amoeboid cells with increased endogenous RASSF1C expression, induced by dCas9-VP64, have elevated co-expression of both contractile actomyosin and Nanog, compared to cells transfected with a repressive Cas9 derivative, dCas9-SKD, where the pMLCII/Nanog^+ve^ pool was significantly reduced (Fig. 4h and Supp. Fig. 4G). The presented data suggest that RASSF1C expression and its downstream Rho/ROCK/pMLCII pathway upregulation re-wire cells to express cancer stemness markers such as CD133, ALDH1, Nanog and OCT4. Having previously shown that RASSF1C/RhoA drive invasive behavior through the release of EVs (Fig. 3), we next wanted to assess whether the cancer stem cell pool of RASSF1C cells promote invasive behavior via EVs. In order to maximize the RASSF1C^+^ cell pool, we sorted ZsGreen-control and ZsGreen-RASSF1C expressing MDA-MB-231 cells for green^high^expression by FACS, and isolated EVs from the sorted pools. Equal amounts of EVs from control and RASSF1C^+^ cells were then incubated with recipient low invading MCF7 cells and cells were allowed to invade through matrigel. The number of invaded MCF7 cells was significantly higher after incubation with Zsgreen^high^ RASSF1C EVs compared to control EVs (Fig. 4i, Supp. Fig. 4H). Furthermore, RASSF1C-derived EVs also displayed the ability to promote upregulation of stem cell markers in recipient MCF7 cells, as shown in Fig. 4j.

### RASSF1C amoeboid cells induce invasion and metastasis via EV transfer *in vivo*

To further corroborate our *in vitro* results and three-dimensional invasion models and validate their relevance *in vivo*, we next employed the previously validated Cre/LoxP system (Fig. 3) in a murine model, to study tumor growth and metastatic potential. The Cre-LoxP system coupled with intravital imaging allows us to investigate *in vivo* invasive behavior of tumor cells, promoted by transfer of extracellular vesicles. We referred to an already established protocol where a mix of either MDA-MB-231^CFP,pUbC-Cre,Control^ or MDA-MB-231^CFP,pUbC-Cre,RASSF1C^ are injected together with T47D^DsRed^ reporter cells into the mammary glands of mice (Fig. 5a and ref. ^34^). Resulting tumors were collected and frozen samples were sectioned in 20 μm slides. We did not observe a difference in the total T47D^DsRed^ tumor area, both in the presence of MDA-MB-231^CFP;pUbC-Cre;Control^ or MDA-MB-231^CFP;pUbC-Cre;RASSF1C^ cells (Supp. Fig. 5A). However, the area of T47D^eGFP^ in MDA-MB-231^CFP;pUbC-Cre,HA-RASSF1C^;T47D^DsRed^ tumors was significantly bigger compared to MDA-MB-231^CFP;pUbC-Cre,control^;T47D^DsRed^ (Fig. 5b images and left bars), suggesting that transfer of RASSF1C-donor derived EVs in the tumor results in higher recombination events, namely reporter cell switch from T47D^DsRed^ to T47D^eGFP^ (Fig. 5b, zoomed images), in agreement with our *in vitro* findings (Fig. 3a, Supp. Fig. 3B). Furthermore, in line with the oncogenic role described for RASSF1C *in vitro*, the area of CFP^+^MDA-MB-231^CFP;pUbC-Cre,HA-RASSF1C^ bearing tumors was significantly greater compared to control (Fig. 5b, right bars). We next evaluated the migratory potential of cells within MDA-MB-231^CFP;pUbC-Cre;Control^ and MDA-MB-231^CFP;pUbC-Cre;RASSF1C^ bearing tumors by time-lapse intravital microscopy. Randomly selected CFP^+^, eGFP^+^ and DsRed^+^ cells from the same field of view were individually tracked and their migration speed was analyzed. In MDA-MB-231^CFP;pUbC-Cre;HA-RASSF1C^ mixed tumors, RASSF1C CFP^+^ cells showed a higher migration speed compared to cells in control mixed tumors (Fig. 5c, d), suggesting that these cells use amoeboid mode of invasion and are characterized by increased migration speed, in agreement with our observations in Fig. 1j. Furthermore, RASSF1C induced T47D^eGFP+^ reporter cells exhibited the greatest migratory potential compared to unrecombined T47D^DsRed+^ cells or control induced T47D^eGFP+^ cells, once again suggesting that RASSF1C driven transfer of EVs has a functional effect also *in vivo*(Fig. 5e, Supp. Fig. 5B, Video 2 for control and 3, 4 for RASSF1C).

**Figure 5.**
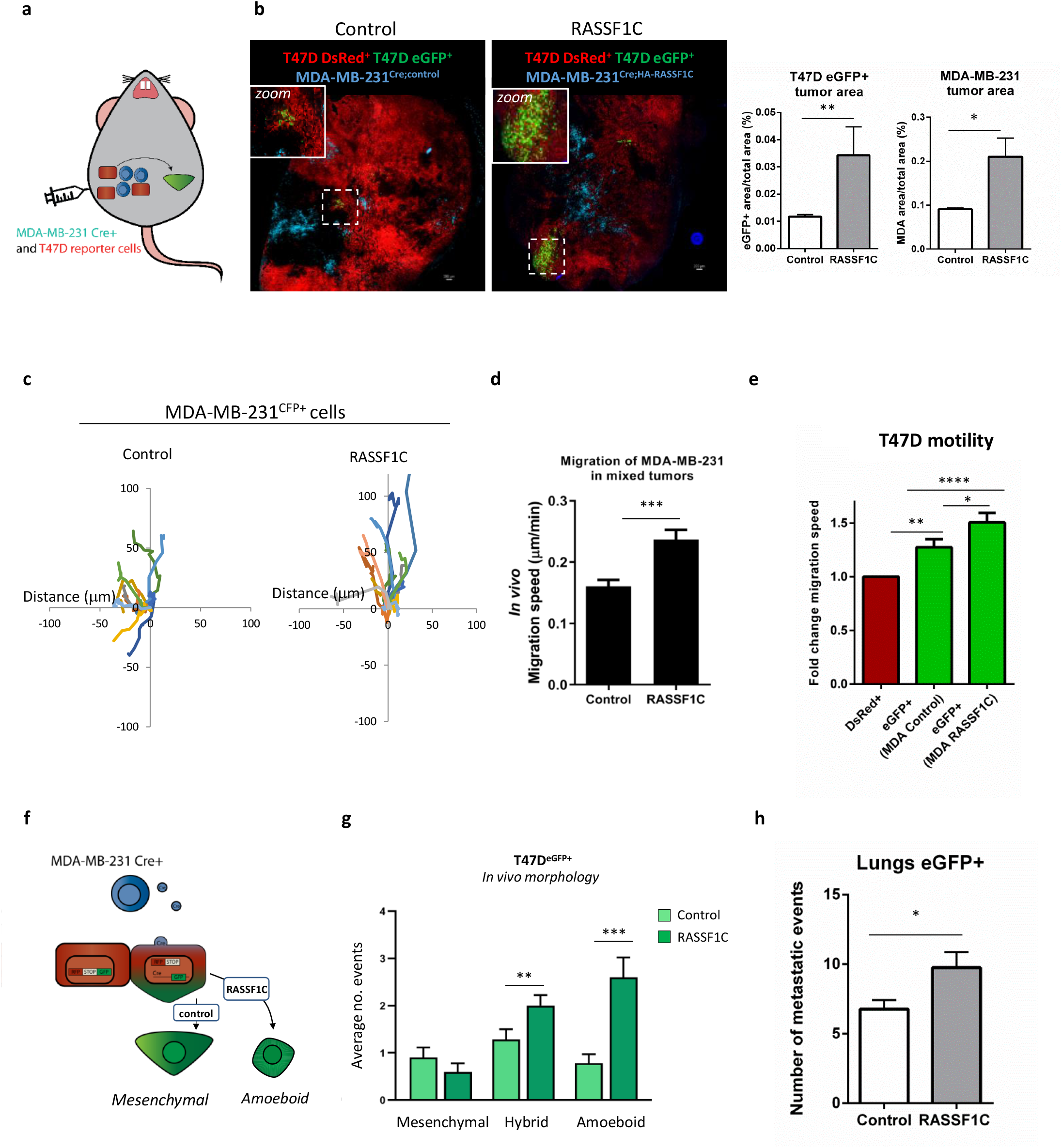
a. Cartoon of in vivo experimental approach. MDA-MB-231^CFP;Cre^ and reported T47D^DsRed^ cells were co-injected in one mammary gland of 10 mice per experiment. b. Left: representative confocal images of mammary tumor formed by the co-injection of T47D^DsRed^ reporter cells with either MDA-MB-231^CFP;Cre;Control^ or MDA-MB-231^CFP;Cre;HA-RASSF1C^. Scale 200 μm. Right: averaged values for all mice for the percent of the eGFP^+^ or CFP^+^ cells as part of the whole tumor. c. Representative 3 hour migration paths of either MDA-MB-231^CFP;Cre;Control^ (Control, left) or MDA-MB-231^CFP;Cre;HA-RASSF1C^ (RASSF1C, right) within a single intravital imaging field of tumors co-injected with T47D^DsRed^ reporter cells. d. *In vivo* migration speed of MDA-MB-231^CFP;Cre;Control^ or MDA-MB-231^CFP;Cre;HA-RASSF1C^ in a mixed tumor with T47D^DsRed^ cells. Values are an average of the totals from 5 mice each, measured from intravital time-lapse images. e. The normalized *in vivo* motility of T47D^eGFP^ and T47D^DsRed^ reporter cells in mixed tumors. A Non-parametric Mann-Whitney U test was used to derive statistical significance. f. Cartoon representing the Cre-LoxP model exploited to follow up amoeboid motility in recombined reporter (T47D^eGFP^) cells. g. Quantification of distinct morphology of T47D^eGFP^ in tumors from observed from a total of 30 positions in n=5 mice/group (Control or RASSF1C). Mesenchymal, Amoeboid or Hybrid motility were scored manually as described in Supp. Fig. 1K. h. Quantification of the number of T47D^eGFP^ metastatic events found in the lungs of mice with mammary gland tumors. Whole lung tile images were used for quantification, 6 of 10 μm sections each, 100 μm apart. All data are from n=3 independent experiments. Data are represented as mean ± SEM.* p ≤ 0.05, ** p ≤ 0.01, *** p ≤ 0.001, *** p ≤ 0.0001.

Using manual tracking we observed that *in vivo* RASSF1C driven eGFP^+^ cells also appear to adopt rounded morphology, typical of amoeboid mode (Fig. 5f, g, Video 2 for control and 3, 4 for RASSF1C), similarly to what we observed for MDA-MB-231^CFP;pUbC-Cre;RASSF1C^ tumors, analyzed in Supp. Fig. 1K. Finally, we analyzed the ability of T47D^eGFP+^ reporter cells to colonize the lungs. A higher number of metastatic events was apparent in T47D^eGFP+^ cells deriving from the MDA-MB-231^CFP;pUbC-Cre,HA-RASSF1C^;T47D^DsRed^ tumors (Fig. 5h, Supp. Fig. 5C for representative images), whereas similar numbers of unrecombined T47^DsRed^ cells migrated to the lungs (Supp. Fig. 5D).

The presented data supports the hypothesis that RASSF1C cells (MDA-MB-231 in our model) adopt amoeboid morphology also in *in vivo* settings and that RASSF1C-mediated EV transfer occurs also *in vivo* and modulates the fate of less invasive cells (T47D in our model), leading to mesenchymal-amoeboid transition, accompanied by increased local motility and distal metastatic events.

### Methylation signature of *RASSF1* gene as a prognostic biomarker in breast cancer patients

Hypermethylation of the *RASSF1* gene has been widely associated with poor prognostic outcome in breast cancer patients ^45^. Specific methylation of the *RASSF1-1α* promoter and loss of the RASSF1A isoform is a frequent event in many tumor types, and offers closer association with cancer progression and poor overall survival ^19,46,47^. However, the impact of *RASSF1-2γ* promoter methylation and expression of RASSF1C, has not been previously taken into consideration (Fig. 6a). Individual *RASSF1* CpG site analysis allows separation of epigenetic silencing at *RASSF1-1α* from *RASSF1-2γ* and therefore study of the differential contribution of the tumor suppressor RASSF1A from the oncogenic RASSF1C isoform to disease progression (Fig. 6a, Supp. Fig. 6A). To determine whether RASSF1C-induced amoeboid invasion and tumor progression may be clinically relevant, we interrogated a TCGA cohort of 732 stage I-IV breast cancer patients, with a particular focus on the early stages (I and II). The TCGA database incorporates tumor methylation signal from 52 CpG sites across the *RASSF1* gene, determined using an Illumina HM450K BeadChip.

**Figure 6.**
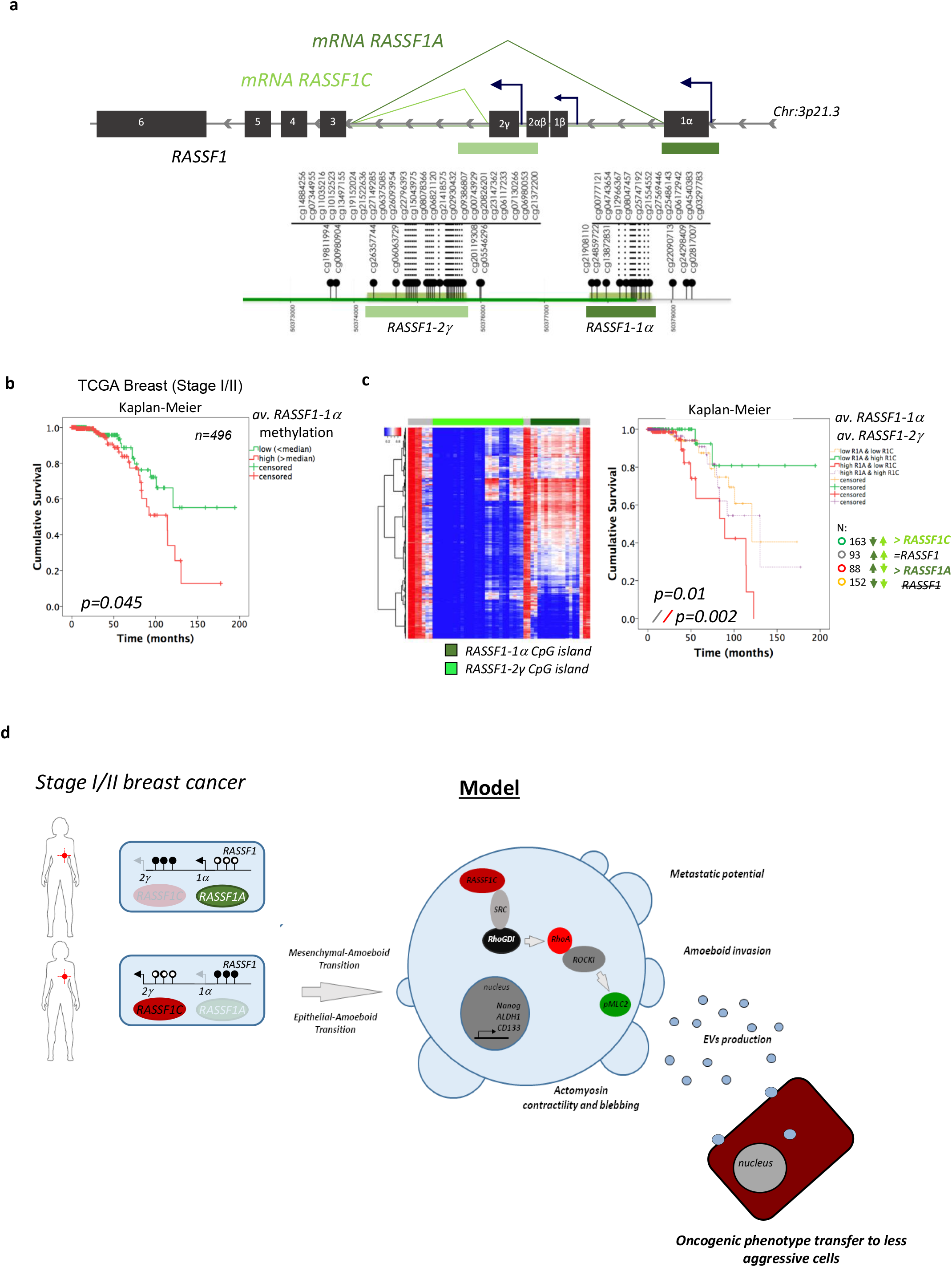
a. Schematic representation of the *RASSF1* gene showing the domains in *RASSF1A* and *C*. Bars shaded dark and light green denote the CpG islands associated with *RASSF1A* and *RASSF1C* respectively. b. Kaplan-Meier plots of overall survival of 496 stage I and II breast cancer patients with different levels of promoter/CpG island methylation of *RASSF1A*, previously reported to be associated with outcome in breast cancer patients. P values were obtained from a log-rank test. c. Left: heatmap depicting methylation pattern across *RASSF1* gene. Right: Kaplan-Meyer survival curves depicting interaction of *RASSF1A* and *RASSF1C* methylation to predict the outcome of early stage breast cancer patients. P values from a log-rank test: yellow, low RASSF1A and low RASSF1C; green, low RASSF1A and high RASSF1C; red, high RASSF1A and low RASSF1C; grey, high RASSF1A and high RASSF1C. d. Model representing how RASSF1C expression in RASSF1A-silenced breast cancer cells (MDA-MB-231) activates a signalling cascade involving SRC kinase, RhoGDI and culminates in RhoA activation and generation of amoeboid contractility and cancer stemness features. RASSF1C-driven amoeboid cancer stem cells contribute to alteration of the tumor microenvironment by rendering recipient cells (MCF7, T47D) more invasive and able to upregulate stem cell markers, due to EV-mediated communication.

We first performed a survival analysis of stage I and II breast cancer patients only considering RASSF1A-1*α* methylation pattern (Fig. 6b). The results support the hypothesis that high methylation of *RASSF1-1α*, indicating reduced RASSF1A expression, is moderately associated with poor breast cancer outcome (p=0.045, log-rank test) (Fig. 6b). Strikingly, *RASSF1-2γ*, the RASSF1C promoter, also demonstrates a spectrum of low to high methylation, suggesting that methylation of *RASSF1-2γ* CpGs may effect clinical outcome (Fig. 6c, heatmap on the left). In general, breast cancer patients that were alive at follow up had lower median methylation of *RASSF1-1α*, in agreement with the tumor suppressor role of RASSF1A, than those stage I and II patients that had died of breast cancer (Supp. Fig. 6B). In line with our data supporting a pro-oncogenic role for RASSF1C, deceased patients had less median methylation of *RASSF1-2γ*, implying that expression from this promoter was active in these tumors (Supp. Fig. 6B). Intriguingly, increased methylation of *RASSF-1α* (reduced RASSF1A expression) has greater significance in stage I tumors (p=0.000001), whereas reduced methylation *RASSF1-2γ*(increased RASSF1C expression) has more prognostic influence in late stage tumors (p=7×10^−6^ Mann-Whitney) (Supp. Fig. 6B table). This supports the possibility of an epigenetic switch in *RASSF1* promoter usage from stage I/II to stage III/IV tumors, leading to RASSF1C expression and RASSF1A silencing ^17^. To determine whether *RASSF1-2γ* methylation status influences stage I and II tumors, we next divided stage I and II breast cancer patients into four groups based on their methylation status, namely *R1-1α^Low^:R1-2γ^Low^, R1-1α^Low^:R1-2γ^High^*, *R1-1α^High^:R1-2γ^Low^*, and *R1-1α^High^:R1-2γ^high^* and repeated the survival analysis from Fig. 6b. In line with the above results, *RASSF1-2γ* methylation (indicating loss of RASSF1C expression) further defines risk groups associated with *RASSF1-1α* methylation and loss of RASSF1A (p=0.01, log-rank test, Fig. 6c Kaplan-Meyer curves). Specifically, the extreme survival groups were those with divergent methylation patterns that would support either RASSF1A (5-yr survival ~ 90% and 10-yr survival ~80%) or RASSF1C expression, (5-yr survival ~60% and 10-yr survival ~10% (p= 0.002, log-rank test, Fig. 6c). Patients with convergent *RASSF1* methylation patterns *(R1-1α^Low^:R1-2γ^Low^* or *R1-1α^High^:R1-2γ^High^)* formed a group with intermediate survival. This indicates that patients with RASSF1C expressing tumors were at over 6-fold higher risk of death than patients likely to express RASSF1A (p=0.02, Supp. Fig. 6C). Intriguingly, the inclusion of intermediate prognosis patients with good prognosis patients offers a significant stratification of patients with different outcomes (data not shown, p=0.002, log-rank test, RR=2.68, p=0.018, Cox multivariate analysis). Notably, no prognostic power for *RASSF1* methylation was seen in n=183 advanced stage breast cancer patients from the same dataset (data not shown).

Taken together, our findings suggest that assessment of the methylation status of both RASSF1-*1α* and *RASSF1-2γ* promoters can be a refined tool for breast cancer patient stratification and concurs to provide a more sensitive prognostic signature than the use of RASSF1-*1α* methylation alone.

## Discussion

The mechanisms underlying metastatic dissemination are largely shaped by the ability of tumor cells to interact with the extracellular environment ^48^. This is achieved by various mechanisms, by direct contact between cells in the tumor tissue, by transfer of secreted molecules or, as more recently described, via secretion of extracellular vesicles (EVs) ^49^. EVs can be internalized by a recipient cell, through membrane fusion or endocytosis processes, and can thereby alter the host cell’s fate ^50^. Numerous *in vitro* and *in vivo* studies have shown that vesicle-mediated exchange of cargo between cancer cells promote cell migration, invasion ^8,51^ and metastatic events ^9,52,53^

Oncogenes have been described as key players in the secretion of pro-invasive EVs. The Rho family member RhoA, one of the major players in amoeboid cell migration, has been previously linked to EV shedding in cancer cells ^11,12^. Here, we describe for the first time activation of the Rho/ROCK/pMLCII pathway through expression of the RASSF1C oncogene, subsequent activation of the SRC kinase and inhibitory phosphorylation of the Rho GTPase inhibitor, RhoGDI.

RASSF1C-driven pathway activation leads to morphological changes in mesenchymal and epithelial cells (Fig. 1a, b) that can be referred to as mesenchymal and epithelial to amoeboid transition, resulting in cytoskeleton reorganization, higher cell invasion (Fig. 1c, g, Supp. Fig. 1F) and *in vivo* dissemination (Fig. 1i). Furthermore, RASSF1C oncogenic function results in secretion of greater numbers of EVs, a process likely to be mediated by RhoA (Fig. 3b, d).

Building on previous work reported by our group on the interplay of the two major *RASSF1* isoforms, RASSF1C and RASSF1A, with the SRC kinase family ^17^, here we report a novel function of RASSF1C/SRC in eliciting amoeboid type of motility. SRC has been previously described to modulate RhoA activity through binding and phosphorylation of the Rho GDP dissociation inhibitor, RhoGDI ^21^. Herein we show that, in the presence of RASSF1C, SRC and RhoGDI precipitate in the same complex, and we suggest that the observed upregulation of Rho/ROCK/pMLCII pathway (Fig. 2) occurs due to failed Rho inhibition by RhoGDI. Importantly, two point mutations generated in the Ras Association (RA) domain of RASSF1C (a domain shared with the tumor suppressor RASSF1A), fail to activate Rho and promote amoeboid transition (Fig. 2f-h). The data suggest that the RA domain, previously shown to be important in RASSF1A direct inhibition of RhoA activity ^14^, may also contribute to RASSF1C activation of RhoA.

In agreement with the oncogenic role of RASSF1C ^19,54^, here we also report evidence that RASSF1C amoeboid cells release EVs which transfer the invasive phenotype to less invasive cells (Fig. 3e-g). RASSF1C-derived EVs fall in the small EVs category (Fig. 3b, Supp. Fig. 3H) and their secretion appears to be sustained by RhoA (Fig. 3d, f, g). Interestingly, analysis of the RASSF1C EV proteome did not reveal differential enrichment of proteins (Supp. Fig. 3D), therefore suggesting that the oncogenic function carried out by RASSF1C EVs could be ascribed to the presence of a specific EV subpopulation, enriched in markers such as CD81, CD63, β-catenin, PDCD6 (as enrichment of these markers was observed in Fig. 3c), or to other types of cargo, such as RNAs or lipids ^34^.

The importance of our *in vitro* results has been translated to *in vivo* settings, where we were able to validate the key role of RASSF1C in promoting tumor progression through EV transfer and amoeboid motility (Fig. 5b, d, e). Remarkably, EVs act both locally and via systemic delivery to distal sites in mice, as previously observed (Fig. 5h and ref. ^34^).

Furthermore, we highlight the relevance of the described biological mechanisms by providing a direct clinical correlation between *RASSF1C* methylation pattern and survival in breast cancer patients (Fig. 6c, Supp. Fig. 6, C).

Heterogeneity of tumors suggests that multiple individual cellular phenotypes exist that can both impart resistance to therapy and have variable metastatic potential ^55^. Moreover, cellular plasticity is now appreciated to allow dynamic switching between phenotypes, further increasing tumor heterogeneity. We previously demonstrated that physiological expression of the tumor suppressor RASSF1A upon stem cell differentiation contributes to cell lineage determination ^56^ and that, conversely, epigenetic silencing of *RASSF1A* in cancer results in tumor microenvironment remodeling and display of cancer stemness properties^57^

RhoA activity has independently been shown to promote stemness via contractility, by inducing nuclear localization of the Hippo pathway mediator YAP1 ^58^ and transcriptional regulation of OCT4 ^59^.

Using both over-expression experiments and epigenetic editing, here we show that RASSF1C drives expression of cancer stemness features in amoeboid invasive cells (Fig. 4b, h) and upregulates a panel of cancer stem cell markers, such as ALDH1, CD133 and Nanog (Fig. 4a, c, d). Furthermore, we show that, as for other RASSF1C-initiated processes, upregulation of stemness markers such as Nanog relies on RhoA activation (Fig. 4e, f, Supp. Fig. 4B-D) and that the same process can be readily initiated by RASSF1C EVs in target cells (Fig. 4j).

Our previous results point at a role for RASSF1C in modulating activity of the Hippo pathway effector YAP1, in the absence of RASSF1A ^17^. Future directions may include exploring the potential link between RASSF1C/RhoA axis and YAP1 function in modulating stemness.

Taken together, our results show for the first time amoeboid motility related to SRC-mediated activation of the Rho/ROCK/pMLCII pathway via the RASSF1C oncogene, in cancer cells where the tumor suppressor *RASSF1A* is epigenetically silenced (schematic model, Fig. 6d). Our results provide new insight on mechanisms of tumor progression, by highlighting the connection between Rho-driven amoeboid motility, upregulation of cancer stemness features and transfer of EVs able to alter the fate of target cells and tissues. The presented results correlate methylation data to disease prognosis and provide an explanation for poor clinical outcome in breast cancer patients with *RASSF1A* methylation. Furthermore, the data point at the interplay between Rho and RASSF1C/SRC pathways as a driver of cancer stemness and aggressiveness. Targeting these pathways would harness existing therapies by preventing cancer plasticity, a phenomenon at the basis of disease progression and dissemination.

## Materials & Methods

### Cell Culture and Transfection

MDA-MB-231, MCF7 and H1299 cells were purchased from ATCC and maintained in DMEM supplemented with 10 % FBS, 2 mM Glutamine and 100 U/ml Penicillin/Streptomycin (all from Thermo Scientific). Stable MDA-MB-231 Cre^+^ and T47D reporter^+^ cells were made as previously described (for reference: PMID 26658469). T47D cells were cultured in DMEM/F12 supplemented with 10 % FBS, 2 mM Glutamine and 100 U/ml Penicillin/Streptomycin. All cells were maintained at 37 °C, with 5 % CO2 in a humidified incubator. Stable cell lines were kept under constant selection using puromycin (Sigma-Aldrich) or Zeocin (Thermo Scientific).

### Transfections and stable cell lines

All cells undergoing transfection were plated in the presence of complete DMEM (Thermo Scientific), without antibiotics. All transfections, either with plasmid DNA, were done using the Lipofectamine 2000 transfection reagent (Thermo Scientific) and following the manufacturer’s guidelines.

Stable cells lines were created using the retroviral vector pQXCIP (Clontech). The cDNAs of ZsGreen, ZsGreen-RASSF1C or HA-RASSF1C were cloned into the pQXCIP construct. pQXCIP-ZsGreen and pQXCIP-ZsGreen-RASSF1C were used to generate stable MDA-MB-231 and MCF7 cells. Empty pQXCIP or pQXCIP-HA-RASSF1C were used to generate stable MDA-MB-231 Cre^+^ cells.

### SDS-PAGE and Western Blot

Cells were lysed for western blot using Laemmli lysis buffer (2.5 mM Tris-HCl pH6.8, 2 % SDS, supplemented with protease inhibitor) before boiling (100 °C, 10 min). Protein sample concentration was measured using Bradford reagent and absorbance measured using POLARstar OMEGA machine at 595 nm. EVs were lysed in 50 μl RIPA buffer (Cell Signalling) containing 20 mM Tris-HCl (pH 7.5), 150 mM NaCl, 1 mM Na_2_EDTA, 1 mM EGTA, 1 % NP-40, 1 % sodium deoxycholate, 2.5 mM sodium pyrophosphate, 1 mM β-glycerophosphate, 1 mM Na_3_VO_4_, 1 μg/ml leupeptin. The samples were then boiled at 100 °C for 10 min and 5 μl of each sample was used for protein quantification using the MicroBCA assay (Thermo Scientific) following the manufacturer’s protocol. The absorbance was then measured at wavelength of 562 nm using a plate reader (BMG POLARstar OMEGA). Samples were normalized with 1 X Loading buffer (Thermo Scientific), so that equal loading per experiment was achieved. Protein samples were loaded onto a gel and separated by SDS-PAGE using NuPAGE^®^ pre-cast gels (10 % or 4-12 %) (Thermo Scientific). Protein was transferred onto PVDF membrane and blocked and incubated with primary antibody in 5 % non-fat milk or BSA diluted in PBS-Tween 20. Secondary antibodies were always incubated in 5 % non-fat Milk. Membranes were covered in ECL solutions from Thermo Scientific, Millipore or GE Healthcare prior to exposure to film (Fujifilm) and developed in a XoGraph developer.

### 2D Immunofluorescence staining

Cells, plated onto glass coverslips (Fisher), were washed twice with PBS and fixed in 4 % (v/v) Paraformaldehyde solution for 15 min after which cells were permeabilized with 0.2 % (v/v) Triton-X solution and blocked with 0.2 % (v/v) Fish Skin Gelatin (FSG) (Sigma–Aldrich) for 1 hr at RT. Cells were then incubated overnight at 4 °C with primary antibody at 1 to 100 dilution. Coverslips were washed 3 times with PBS before the Alexa fluor secondary antibody (1 in 500) (Fisher Scientific) was added and the cells incubated for a further 1 hr at RT. Coverslips were washed with PBS and mounted onto microscope slides (Fisher) using Prolong® Gold antifade reagent with DAPI (Thermo Scientific). Fluorescent microscopy was done using a Nikon Ti-E or a Nikon 2000TE microscope using the NIS elements software (Nikon), version 4.2. Confocal microscopy was done using a Zeiss LSM780 with Zen 2011 program.

### 3D Immunofluorescence staining

Multicellullar aggregates were generated for 24 hours by using the hanging drop method ^60^. Spheroids were washed and either seeded on Matrigel matrix for another 48 hours in complete medium or mixed with collagen matrix. For 3D spheroids in three-dimensional collagen staining, collagen-spheroids gels were washed with PBS, fixed with 4 % PFA for 30 minutes and crosslinked in sodium azide solution overnight. After washing with PBS, spheroids in collagen gels were incubated in primary antibodies diluted in 0.2 % Triton and 10 % NGS overnight, extensively washed with PBS and incubated in Alexa Fluor secondary antibodies diluted in 0.2 % Triton and 10 % NGS solution, overnight in 4 °C. Spheroids-collagen gels were washed with PBS and mounted with DAPI. 3D spheroids cultured on Matrigel matrix were washed with PBS, fixed with 4 % PFA for 1 hour and 3 times washed with PBS:Glycine solution (NaCl, Na_2_HPO_4_, NaH_2_PO_4_, glycine, pH 7.4). Spheroids were permeabilized with 0.5 % Triton X-100 for 10 minutes, washed 3 times with IF solution (NaCl, Na_2_HPO_4_, NaH_2_PO_4_, NaN_3_, BSA, Triton X-100 and Tween solution, pH 7.4) and blocked for 2 hours in 10 % goat serum in IF solution. Spheroids on Matrigel were then incubated with primary antibodies diluted in 10 % goat serum and IF solution for 2 hours, washed 3 times with IF solution and incubated in secondary Alexafluor antibodies diluted in IF solution for 1 hour. Spheroids were then washed with IF solution and mounted with DAPI. Images were captured by using Nikon 10x/0.30 Ph1 objectives.

### In-Gel Gelatin Zymography

MDA-MD-231 cells were counted and 2 × 10^5^ were plated into 24-well plate. After seeded, cells were washed with PBS and incubated in 300 μl of serum-free medium for 72 h. Aliquots of the conditioned medium were mixed with 4 X sample buffer and loaded for zymography on a 10 % SDS-PAGE gel containing 1 mg/ml gelatin as previously published ^61^. Briefly, gel containing metalloproteases was washed for 1 h in 50 mmol/L Tris-HCl (pH 7.5), 0.1 mol/L NaCl, and 2.5 % Triton X-100 and then incubated at 37 °C in 50 mmol/L Tris-HCl (pH 7.5), 10 mmol/L CaCl_2_, and 0.02 % sodium azide overnight. The gels were stained with Coomassie blue then destained in 7 % acetic acid/5 % methanol and imaged for gelatin degradation area.

### Reverse Transcriptase and Quantitative Real Time PCR (qRT-PCR)

Reverse Transcriptase PCR and Quantitative Real-Time PCR were prepared using the Power SYBR Green Cells-to-Ct Kit (Thermo Scientific) protocol. Cells (2 x 10^5^/well) were plated onto 6 well plates and 2 days later 5 x 10^4^ cells were used for cDNA preparation. Samples were run on Applied Biosystems 7500 Fast real-time thermal cycler (Applied Biosystems). The protocol used was Holding Step (1 X cycle: 95 °C 10 min), Cycling Step (50 X cycle: 95 °C 15 sec, 60 °C 1 min) and Melt Curve Step (1 X cycle: 95 °C 15 sec, 60 °C 1 min, 95 °C 30 sec, 60 °C 15 sec). 18S or GAPDH were used as internal controls for all the experiments. The calculation of the ΔCt was done on Microsoft Excel.

### EV Isolation and Treatment of cells

Cells (1 x 10^7^ cells/condition) were grown for the indicated times, up to 3 days, in DMEM supplemented with 0.1 % (v/v) FBS (depleted of bovine EVs by overnight centrifugation at 100,000*xg*), 2 mM Glutamine and 100 U/ml Pen/Strep. Conditioned medium was collected and EVs were isolated by sequential ultracentrifugation at *2000xg* for 30 min, 10,000*xg* for 40 min, 100,000*xg* for 3 h in an Optima XPN-80 (Beckman Coulter) ultracentrifuge using UltraClear Thinwall tubes. The EVs were washed once in 1 ml of PBS and purified by centrifugation at 100,000*xg* for 80 min in an Optima MAX-XP Ultracentrifuge (Beckman Coulter). EV protein concentration was measured using MicroBCA assay (Thermo Scientific) to ensure equal amounts were added to cells. Unless stated otherwise, 20 ng/μl of EVs were used per well. When stated, prior to addition to cells, EVs were stained using the PKH67 Green Fluorescent Cell Linker Mini Kit (Sigma–Aldrich) following manufacturer’s protocol. Internalization of EVs was assessed by confocal microscopy at a Zeiss LSM780 microscope.

### EV Isolation by Tangential Flow Filtration/Size Exclusion Chromatography

Cells were handled as described under ‘EV Isolation’. EVs were isolated from ~ 80 mL conditioned medium as previously described ^38^. Briefly, the 10,000*xg* supernatant was concentrated down to 1.0 ml with a 10 kDa molecular weight cut-off Vivaflow 50 R tangential flow (TFF) device (Sartorius) and a 10 kDa Amicon Ultra-15 centrifugal filter unit (Merck Millipore, Billerica, Massachusetts, USA). The concentrated supernatant was loaded on a Tricorn 10/300 Sepharose 4 Fast Flow (S4FF) size exclusion chromatography column (GE Healthcare, Buckinghamshire, UK) connected to the ÄKTA prime system (GE Healthcare), and eluted at 0.5 ml/min flow rate using PBS as the eluent. Chromatogram was recorded using absorbance at 280 nm. 1.0 ml fractions were collected, and EV-containing fractions were pooled and concentrated with a 10 kDa molecular weight cut-off Amicon Ultra-4 centrifugal filter unit (Merck Millipore).

### Nanoparticle Tracking Analysis (NTA)

For this analysis 1/5 of the total EV pellet per sample was resuspended in 10 μL PBS (Gibco) and stored at 4°C for up to 7 days. Additional 990 μL PBS were added on the day of analysis and the EV pellet was vigorously resuspended by pipetting and kept on ice, before starting the analysis. Before starting any measurement, NanoSight NS300 (Malvern Panalytical) was washed three times by loading distilled water onto a syringe pump using a 1 mL syringe and pressing the liquid into the flow-cell top plate of the NanoSight. PBS was used to prime the instrument and to control the purity of the diluent (i.e. absence of particulate in the solution or presence of particulate in a concentration lower than detectable level). After priming, 1 mL of sample was carefully loaded on the syringe pump. Every measurement was done automatically, with the aid of the syringe pump and a script for data acquisition was generated on the NTA 3.2 software. Three recordings of 60 min each were automatically taken once each sample was loaded in the chamber and the focus on the particles in solution was adjusted manually. Additionally, every sample was loaded onto the chamber twice and an average of 6 recordings was then used as a final value. NTA 3.2 software was used to generate data output.

### Time Lapse Imaging

MCF7 cells (3 x 10^5^ cells/well) were plated onto 12-well plates. Cells were incubated for 24 h before media was changed to complete DMEM/F-12 media without phenol red. The cells were then imaged for up to 8 h at a rate of 1 picture every 15 min, using a Nikon TE2000 inverted microscope at 20 x magnification at 37 °C and 5 % CO_2_. The collected pictures and videos were analysed using Nikon NIS Elements 4.0 program. Where necessary, still images were used at 0 h and 12 h to quantify the number of rounded, spherical cells per field of view. The diameter was also quantified using Nikon NIS Elements 4.0.

### Fluorescence Assisted Cell Sorting

Fluorescence Assisted Cell Sorting (FACS) of Green^high^ cells was performed as follows. Briefly, cells (1 x 10^6^) grown in 2D were trypsinised and washed in PBS. Cells were resuspended in PBS containing 1 % FBS. Cells were gated and sorted for high green fluorescence on a BD FACSAria III. Sorted cells were re-plated and grown in full medium. Wild type MDA-MB-231 cells were used as control for the set up and the definition of the fluorescence threshold.

### Immunoprecipitation

1 x 10^6^ cells per condition were transfected with the described plasmids, as stated in ‘Transfection methods’. Cells were washed two times with PBS before lysis with immunoprecipitation buffer (20 mM TrisHCl pH 7.4, 1 % NP40, 100 mM NaCl, 0.5 mM EDTA, 1 X protease and 1 X phosphatase inhibitors (Roche), 10 % v/v glycerol). Cells were scraped from the dish and collected into a 1.5 ml microfuge tube and incubated end-over-end at 4 °C for 10 min. Lysates were then cleared by centrifugation (20817*xg*, 20 min, 4 °C). Protein concentration of the lysates was determined by Bradford assay, so that an equal amount of protein was loaded into each immunoprecipitation reaction.

Protein A Dyna-beads (Millipore) were used for immunoprecipitation. All beads were washed 3 times in immunoprecipitation buffer before being aliquoted into 1.5 ml microfuge tubes. Each lysate was pre-cleared with 10 μl Protein A Dyna-beads for 30 min prior to immunoprecipitation. 1 μg of antibody or corresponding amounts of same species IgG (Cell Signalling) were added to 20 μl of protein A Dyna-beads and incubated end-over-end at 4 °C for 30 min before lysate was added to the tubes and incubated end-over-end at 4 °C for 3 h. Samples containing protein A Dyna-beads were washed three times by immobilizing the beads using a magnet, aspirating the supernatant and resuspending the beads. Protein was removed from the antibody by boiling at 100 °C for 5 min in 2 X LDS Sample buffer (Life Technologies). An equal volume of each sample and its corresponding input was loaded onto a gel for electrophoresis and western blot analysis.

### Migration Assay with xCELLigence

160 μl DMEM containing 10 % FBS were added to the bottom wells of xCELLigence Real-Time Cell Analyzer CIM plate (Roche), as chemo-attractant. The top chamber was then attached to the bottom chamber and 50 μl of DMEM without supplements were added to each well. The plate was first incubated for 30 min at 37 °C and then loaded into the xCELLigence analyser and a blank run was run to zero the machine. EVs from described conditions were resuspended in 50 μl sterile PBS (Gibco) and normalized according to BCA assay (Thermo Fisher), in order to equally load 20 or 40 ng/μl of EVs per well. 50,000 cells were resuspended in DMEM without supplements, with or without EVs. Cells were plated into the top chamber and allowed to settle for 30 min at RT before the plate was loaded into the xCELLigence analyzer. Cells were incubated in the machine at 37 °C, 5 % CO2 in a humidified incubator and measurements were taken every 15 min for 36 hr. Normalization was calculated by subtracting the invasion rate of untreated MDA-MB-231 cells to the invasion rate of EV-treated MDA-MB-231 cells.

### 3D Spheroids formation and morphology in 3D matrix

MDA-MB-231 cancer multicellular spheroids were generated by using the hanging drop method ^60^. Briefly, ZsS-Ggreen-control or Zs-Ggreen-RASSF1C transfected cells were detached with 2 mmol/L EDTA, counted, re-suspended in DMEM supplemented with methylcellulose (20 %, Sigma) and GFR Matrigel matrix (1 %, Corning) and incubated as droplets (25 μl) containing 1,000 cells for 24 hours to generate multicellular aggregates. Three dimensional aggregates were washed with medium, and either seeded and cultured on Matrigel matrix for 48 hours or mixed with rat-tail collagen (Serva; 2.0 mg/mL), 10 X PBS, 1M NaOH and complete medium. Collagen gel-spheroids solution was pipetted as a 100 μl drop-matrix suspension into 12-wells, polymerized at 37°C, replaced with medium and cultured for another 24 hours till single cell invasion from spheroids was apparent. Single cell morphology apparent as mesenchymal (fibroblast-like) projections from the ZsGreen transfected spheroids or as amoeboid (rounded) single cells invaded around from ZsGgreen-RASSF1C transfected spheroids was imaged and captured by using Nikon 20x/0.8 Ph1 objective.

### Boyden chamber invasion and Hydrogel invasion assays

MDA-MB-231 or MCF7 cancer cells were pre-treated or not with either 10 μmol/L of ROCK inhibitor Y-27632 for 1 hour or with 2 μg/ml Rho inhibitor C3 for 4 hours. Cells were trypsinized and (1 × 10^5^) were cultured in serum-free medium alone with 20 μmol/L GM6001 metalloprotease inhibitor, or incubated with EVs (20 or 40 ng/μl) in the upper wells (in triplicate) of Transwell Matrigel Boyden chambers (Corning), or seeded in upper wells coated with 50% hydrogel matrix (PuraMatrix, Corning). Cells were allowed to invade toward bottom wells supplemented with 10% FBS medium. After 8 or 18 hours of incubation, invading cells were fixed and stained with Richard-Allan Scientific™ Three-Step Stain (Thermo Fisher Scientific), photographed, and counted manually using Adobe Photoshop software. Images were captured by Nikon Ti90 4x objective.

### Single cell morphology in 3D collagen

To analyze single cell morphology in 3D collagen, cells were trypsinized, washed in with complete medium, counted and (10^5^/ml) cells were mixed with a solution containing 2mg/ml Collagen R (Serva), 10 X PBS, 1M NaOH and complete medium. The suspension of cells was loaded in 96 wells. The gels were polymerized at 37 °C for 30 min and replaced with complete medium. After 24hours cell morphology in 3D collagen was analyzed by using a Nikon Eclipse TE2000 (10x/0.40 Ph1 objective) microscope, counted and classified on the basis of the elongation index. The elongation index was calculated as the length divided by the width. For each condition a minimum of 500 cells were analyzed.

### Immunoblotting and Rho pull-down assay

Confluent cell cultures were washed with cold 1 X PBS and lysed in modified RIPA buffer (50 mM Tris-HCL (pH 7.4), 150 mM NaCL, 5 mM EDTA, 1 % NP40, 1 % sodium deoxycholate, 50 mM NaF and 1 % aprotinin and 0.1 mM Na_3_VO_4_). Rho pull-down assays were performed using GST-rhotekin and Rho-pull down detection kit according to manufacturer’s instructions (Thermo Scientific). The levels of total and active Rho were established by using a Rho-specific antibody. Protein concentration in lysates was determined by BCA assay. For immunoblotting, samples were separates on 12 % SDS-polyacrylamide gels and transferred onto nitrocellulose membrane. Non-specific activity was blocked by 1 X TBS containing 0.05 % Tween and 4 % bovine serum albumin (BSA). Membranes were incubated overnight in Rho primary antibody, washed and then incubated within HRP-conjugated secondary antibody for 1 hour at RT. After extensive washing with TBS-T, blots were developed as previously stated.

### CRISPR

Plasmids containing a mammalian codon-optimized dCas9-VP64 activator (pMLM3705, AddGene #47754) and single-chain gRNA encoding plasmid (MLM3636, AddGene #43860) were gifts from Keith Joung. Plasmids encoding dCas9-only and dCas9-Super Krab Domain (SKD) repressor were gifts from Marianne Rots ^62^. gRNAs were cloned into BsmBI-digested MLM3636 plasmid by ligation of annealed oligonucleotides encoding 20 bp gRNA target sequences and 4 bp overhangs. Four target regions of 20 bps of the RASSF1C promoter were selected to design gRNAs based on close proximity to the transcription start site (TSS) (gRNA1: TTGTGCGCTTGCCCGGACGC; gRNA2: CGGAGCGATGAGGTCATTCC; gRNA3:

GGATCTAGCTCTTGTCTCAT reverse strand; gRNA4: AGTGCGCGTGCGCGGAGCCT reverse strand). For dCas9 experiments, each well of a 12-well plate was transfected with 500 ng of dCas9 construct and 500 ng combined gRNAs. Transfected cells were collected 48 hours post transfection.

### Co-cultures and Transwell experiments

For all co-culture and transwell experiments, complete DMEM culture medium was used. MDA-MB-231^CFP;Cre^ cells were co-cultured with T47D cells for one week prior to imaging in a culture dish. All experiments were done in parallel. For the transwell experiments, transwells with a pore-size of 400 nm (Greiner Bio-one, Frickenhause, Germany) were place in a culture dish with T47D cells for one week prior to imaging. The cells in the culture dish were imaged with a Nikon 2000TE microscope using a 20x objective. During imaging, cells were kept at 37 °C in a humidified atmosphere containing 5 % CO2. A 430/24 nm excitation filter and a 470/40 nm emission filter was used for CFP, a 470/40 nm excitation filter and a 520/40 nm emission filter was used for eGFP, and a 572/35 nm excitation filter and a 640/50 nm emission filter was used for DsRed.

### Mice

NOD scid gamma (NSG) mice (own crossing) were housed under IVC conditions. Mice received food and water ad libitum. All experiments were carried out in accordance with the guidelines of the Animal Welfare Committee of the Royal Netherlands Academy of Arts and Sciences, The Netherlands.

### Injection of tumor cells

One week before injection of tumor cells, mice were ovarectomized and implanted with an oestrogen pellet (0.36 mg/pellet, 60-day release, Innovative Research of America, Sarasota, FL, USA) using a precision trochar, as per the manufacturer’s instructions. Tumor cells were injected in the fourth and/or ninth mammary fat pad of female 8-20 weeks old NSG mice. For MDA-MB-231 tumor development, 5 x 10^5^ MDA-MB-231 cells in sterile PBS were injected. For MDA-MB-231 and T47D mixed tumors, a total of 1 x 10^6^ cells was injected in a ratio of 10:1 Cre^+^:reporter^+^ cells in sterile PBS with 50 % growth factor-reduced Matrigel (BD Biosciences, Franklin Lakes, NJ, USA).

### Tumor and tissue processing

Tumors and tissues were removed from the mice at the end of the experiment and fixed and processed as described before (for reference: PMID 26658469). In short, tissues were fixed in periodate-lysine-paraformaldehyde (PLP) buffer (2.5 ml 4 % PFA + 0.0212 g NaIO4 + 3.75 ml L-Lysine + 3.75 ml P-buffer (pH 7.4)) O/N at 4 °C. The following day, the fixed tumors and tissues were washed twice with P-buffer and placed for at least 6 hours in 30 % sucrose at 4 °C. The tumors and tissues were then embedded in tissue freezing medium (Leica Microsystems, Nussloch, Germany) and stored at −80 °C before cryosectioning.

### Immunostainings and Confocal Microscopy of tissue sections

Tumor or tissue cryosections (15 μm thick when used for subsequent immunostainings) were rehydrated for 10 min in Tris 0.1 M pH 7.4, when indicated used for immunostainings and embedded in Vectashield mounting medium (hard set; Vector Labs, Burlington, Ontario, Canada). When indicated, cryosections were counterstained with 0.1 μg/ml DAPI (Invitrogen Life Technologies, Paisley, UK) to visualize the nuclei. Images were acquired using Nikon TiE microscope equipped with 40x dry objectives. DAPI and CFP were excited with a UV 405 nm laser, and emission was collected at 415-455 nm for DAPI and 455-495 nm for CFP. eGFP was excited with an argon ion laser at 488 nm and emission was collected at 490-515 nm. DsRed was excited with a 561 nm laser and emission was collected at 570-620 nm. At least 6 frozen sections per mouse were analyzed. Area and number of primary tumors and metastatic events to the lungs were quantified using Nikon NIS software.

### Intravital Imaging

Mice were sedated using isoflurane inhalation anaesthesia (1.5 % to 2 % isoflurane/O_2_ mixture). The imaging site was surgically exposed, and the mouse was placed with its head in a facemask within a custom designed imaging box. The isoflurane was introduced through the facemask, and ventilated by an outlet on the other side of the box. The imaging box and microscope were kept at 36.5 °C by a climate chamber that surrounds the whole stage of the microscope including the objectives. Imaging was performed on an inverted Leica TCS SP5 AOBS or Leica SP8 multi-photon microscope (Mannheim, Germany) with a chameleon Ti:Sapphire pumped Optical Parametric Oscillator (Coherent Inc. Santa Clare, CA, USA). The microscope is equipped with four non-descanned detectors: NDD1 (<455 nm), NDD2 (455-490 nm), NDD3 (500-550 nm), NDD4 (560-650 nm). CFP was excited with a wavelength of 840 nm, and GFP and DsRed were excited with a wavelength of 960 nm. CFP was detected in NND2, GFP was detected in NDD3, and DsRed in NDD4. All images were in 12 bit and acquired with a 25x (HCX IRAPO N.A. 0.95 WD 2.5 mm) water objective. All pictures were processed using ImageJ software or Imaris 8.2 (Bitplane); pictures were converted to an 8 bit RGB, smoothed (if necessary), cropped (if necessary), rotated (if necessary) and contrasted linearly. At the beginning of each movie, a random DsRed^+^ cell close to an eGFP^+^ was selected. The XY position was determined over time and the displacement and track distance was calculated by Excel (Microsoft).

### In silico analysis

Breast cancer data, including clinical and methylation was downloaded from TCGA database (Broad Institute TCGA Genome Data Analysis Center (2015): Analysis-ready standardized TCGA data from Broad GDAC Firehose stddata 2014_07_15 run. Broad Institute of MIT and Harvard. Dataset.

https://doi.org/10.7908/C1TQ60P0). A total of 732 patients had HM450K tumor methylation data. Clinical data was available for 695 patients, 687 of which were females, with an average age at diagnosis 58 years (+/− 13 years). Tumor stage was determined for all 687 female patients, whereby 114 patients were stage I at primary diagnosis, stage II-384 patients, stage III-176 patients and stage IV-7 patients. Oestrogen receptor (ER) status was determined for 656 primary tumors, whereby 498 were ER positive and 150 were ER negative. Male patients were excluded from analyses. Molecular subtype (PAM50) was available for 209 tumors, whereby 107 had Luminal A subtype, 43 Luminal B, 40 Basal, 14 Her2+ and 5 normal-like.

The CpG island methylation was calculated as the average methylation of CpG sites located within defined CpG island at Genome Browser (http://genome-euro.ucsc.edu/index.html). The median values were calculated for all 687 female patients.

### Statistics

For all *in vitro* experiments, unless stated otherwise in figure legends, statistical analysis was carried out using Student’s T-test, calculated on Microsoft Excel or Prism 6.0 (Graphpad). The Standard Error Mean (SEM), used for error bars, was calculated by the division of the standard deviation by the square root of the number of measurements made. For *in silico* analysis statistics were performed using SPSS 24.0 and R version 3.3.2. The heatmaps were created using the ggplot package. A Mann-Whitney-Wilcoxon Test was used to compare two different sample populations. Statistical significance was regarded as P < 0.05.

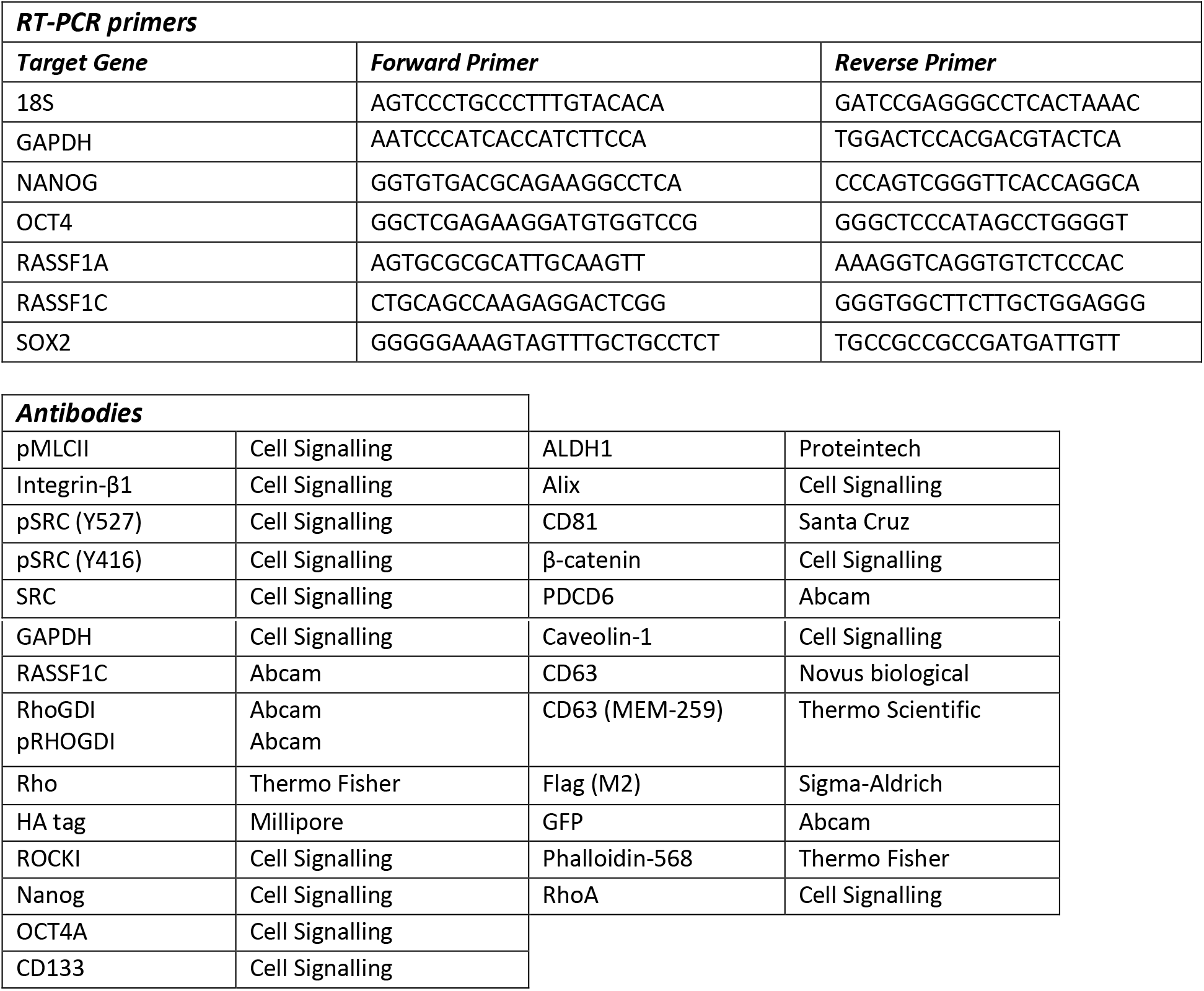

## Author Contributions

DP, MLT and NV performed all *in vitro* work; NV and SSt performed all *in vivo* work and statistical measurements under guidance of JvR; ME and DCR employed the CRISPR dCas9 experiments under the guidance of MGR. EW and MLT performed SEC isolation under the supervision of MJAW. AMG performed *in silico* analysis on TCGA retrieved data with the help of CK. AvK performed all proteomic experiments, EON designed and supervised all experiments, conceptualized the story and wrote the manuscript with the help of DP and MLT.

## Acknowledgements

Authors would like to specifically thank A.L. Harris for critical comments and V. Sanz-Moreno for sharing data prior to publication, J. Brabek and D. Rosel, Charles University in Prague, and P. Vader, University Medical Centre Utrecht, for their input on the manuscript. This work was funded by the MRC, Oxford Cancer Research Centre, Cancer Research UK research travel award, EU cost action CM1406 and Cancer Research UK A19277.

## Supplementary Figure Legends

**Supplementary Figure 1.**
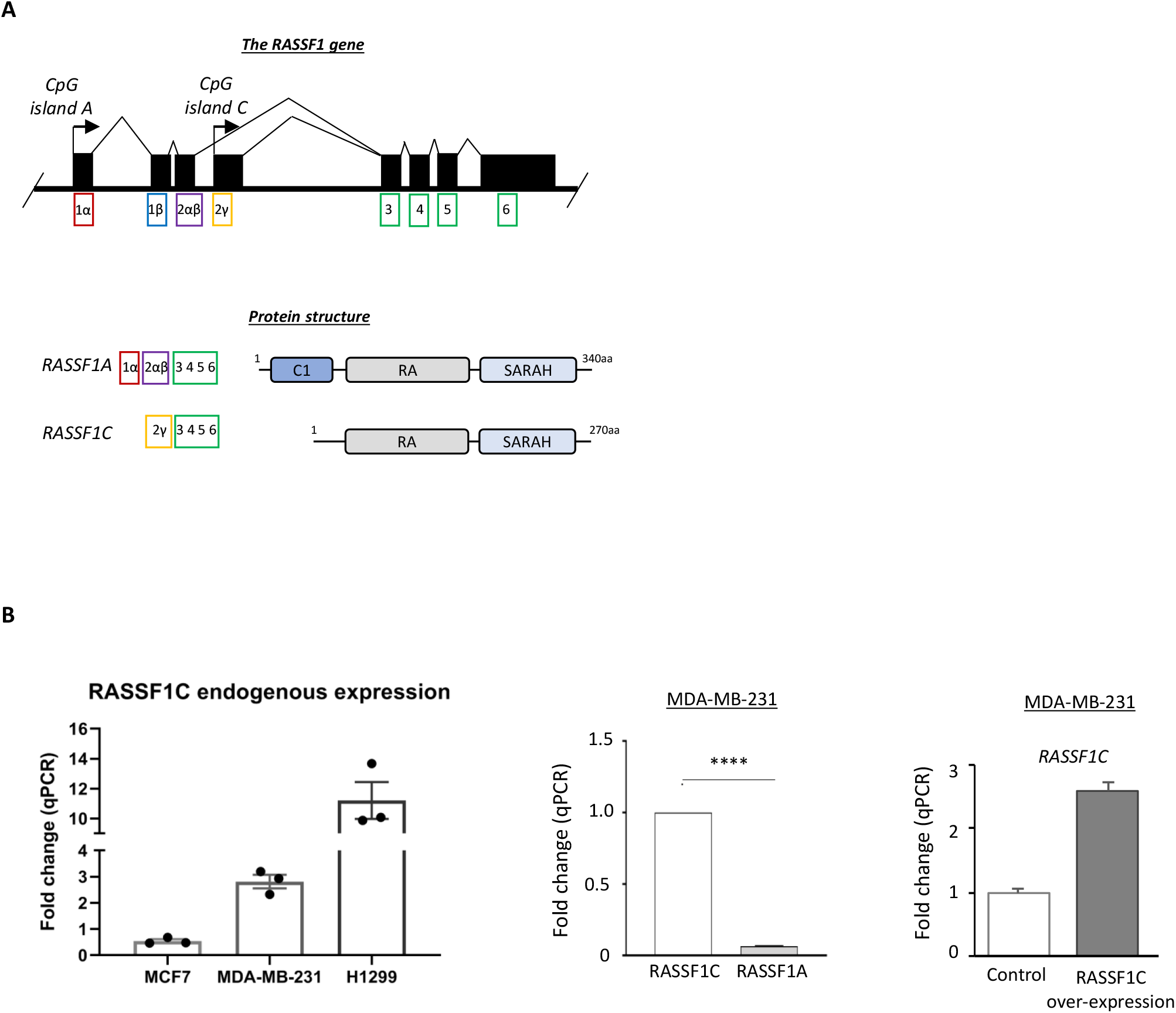
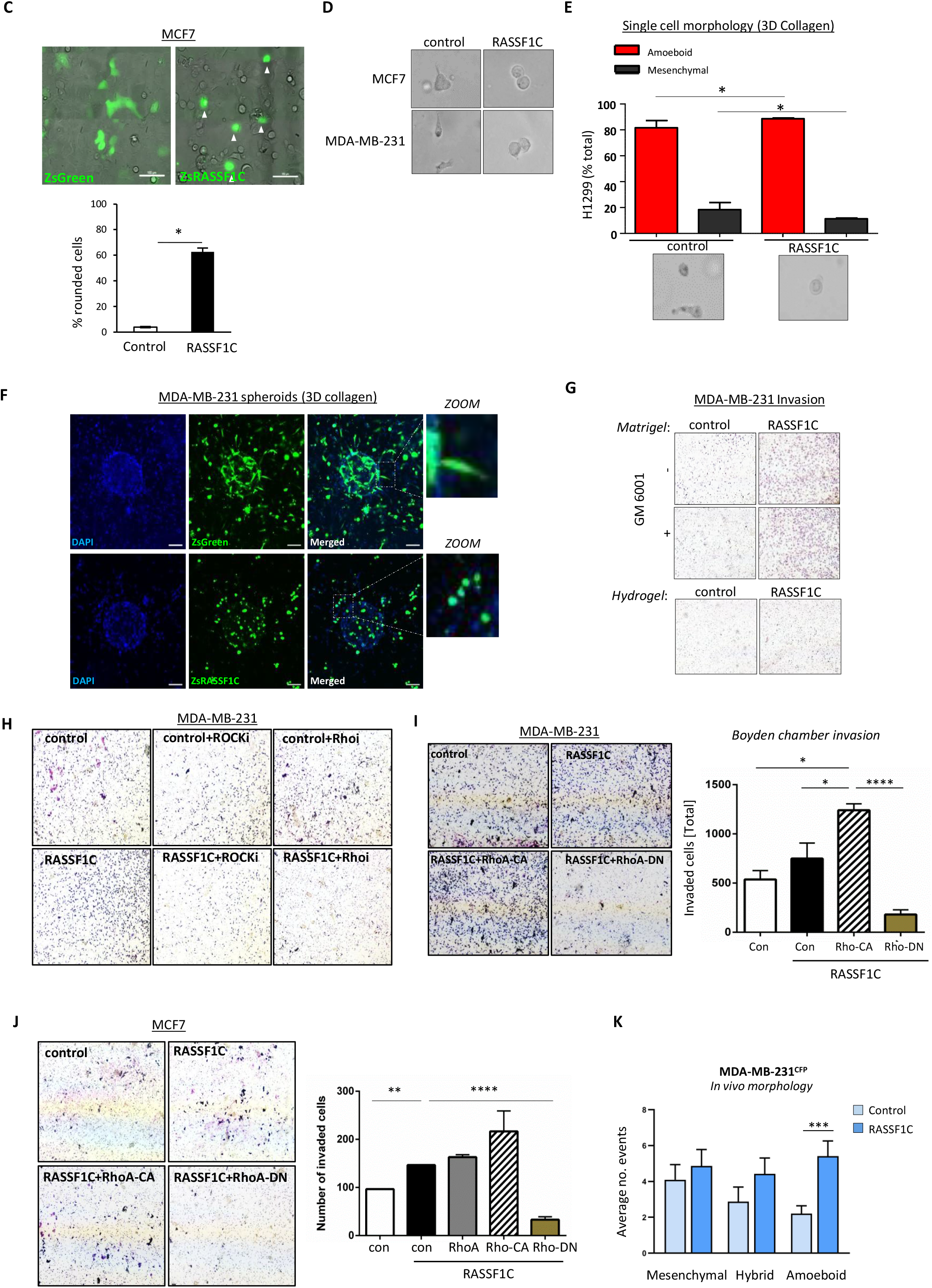
A. Schematic representation of the *RASSF1* gene locus (top) and protein domains of the two main *RASSF1* isoforms (bottom). Top: black boxes represent exons, black arrows represent CpG island promoters. Bottom, RASSF1A transcript contains six exons (1α, 2αβ, 3, 4, 5 and 6) and is translated to a protein with 340 aa. The RASSF1C variant is transcribed from an intragenic CpG island and consists of five exons (2γ, 3, 4, 5 and 6) and is translated to a protein of 270 aa. Bottom: at the protein level, RASSF1A and RASSF1C both encode for a RA (Ras Association) domain and a C-terminal SARAH (Sav/RASSF/Hpo) domain. RASSF1C however lacks the C1 (protein C kinase conserved) region at the N-terminal, which is present in RASSF1A. B. Left: fold change of qRT-PCR for endogenous RASSF1C expression in the cells lines used in this study is shown. Middle: fold change of qRT-PCR for RASSF1C and RASSF1A mRNA in MDA-MB-231 cells are shown. Right: MDA-MB-231 cells were transfected with a Control (pcDNA3) or FLAG-RASSF1C plasmid and mRNA levels of RASSF1C were analysed via qRT-PCR. Data show a 2.5-fold increase in RASSF1C mRNA upon overexpression. C. Top: images taken 48 h after MCF7 cells were transfected with ZsGreen or ZsGreen-RASSF1C (ZsRASSF1C) plasmids. Scale bars 100 μm. Bottom: quantification of ZsGreen^+^ cells with rounded morphology as percentage of the total number of ZsGreen^+^ cells. D. Related to Fig. 1b, representative images of single cell morphology of MCF7 and MDA-MB-231 cell lines, expressing or a ZsGreen or ZsGreen-RASSF1C construct, cultured in 3D rat tail collagen I and analysed for their amoeboid or mesenchymal (fibroblast-like) morphology, 24 hours after seeding in 3D matrix. E. Quantification (top) and representative images (bottom) showing single cell morphology in 3D-collagen matrix in H1299 cells, 12 hours after being transfected with ZsGreen (Control) or ZsRASSF1C (RASSF1C) plasmids. F. Immunofluorescent images of ZsGreen or ZsRASSF1C expressing MDA-MB-231 spheroids cultured in 3D rat tail collagen I. Zoom images showing detail of single cells morphology adapted during invasion from 3D spheroids. Scale 100 μm. G. Related to Fig. 1f, representative images of 3D Matrigel Boyden chamber invasion with or without metalloproteases inhibitor GM6001 and 3D Hydrogel invasion of MDA-MB-231 control cells or cells overexpressing RASSF1C. H. Related to Fig. 1g, representative images of Boyden chamber invasion in 3D Matrigel of MDA-MB-231 cells, transfected with ZsGreen or ZsRASSF1C and treated with inhibitors against ROCK (Y-27632, 10 μmol/L) or Rho (C3, 2 μg/ml). I. Representative images and quantification of Boyden chamber invasion in 3D matrigel of MDA-MB-231 cells, transfected with ZsGreen-Control or ZsGreen-RASSF1C and RhoA catalytically active (CA) or dominant negative (DN) plasmids. J. Representative images and quantification of Boyden chamber invasion in 3D matrigel of MCF7 cells, transfected with ZsGreen-Control or ZsGreen-RASSF1C and RhoA catalytically active (CA) or dominant negative (DN) plasmids. K. Quantification of CFP^+^ MDA-MB-231 cells displaying different types of morphology *in vivo*, expressed as average number of cells, observed from a total of 30 positions in n=5 mice/group (Control or RASSF1C). Mesenchymal, amoeboid and hybrid morphology were scored manually based on diameter and circularity (distance between the two furthest points). The higher distance between cell edges was defined as mesenchymal morphology, while the lower distance was considered amoeboid morphology. Data are represented as mean ± SEM.* p ≤ 0.05, ** p ≤ 0.01, *** p ≤ 0.001.

**Supplementary Figure 2.**
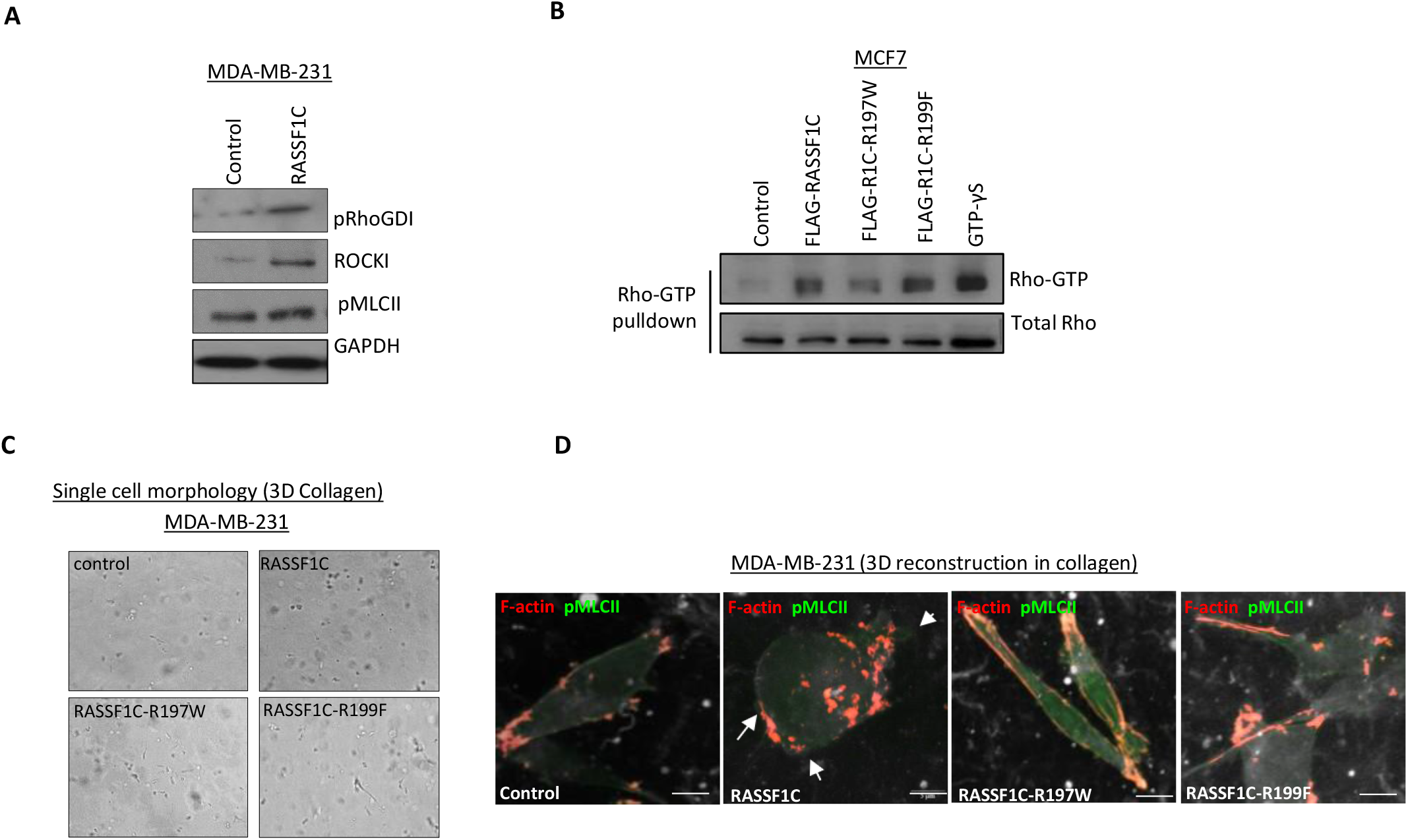
A. Western blot analysis of protein expression in MDA-MB-231 cells transiently transfected with an empty vector or a RASSF1C construct. GAPDH was used as a loading control. B. Active Rho-GTP pull down assay from MCF7 cells transfected with Control, FLAG-RASSF1C, FLAG-RASSF1C R197W or FLAG-RASSF1C R199F plasmids. C. Related to Fig. 2h, representative images, from which quantification in 2h was derived, of single cell morphology assay in 3D-collagen indicating mesenchymal-amoeboid transition of MDA-MB-231 cells when RASSF1C or mutant derivatives are expressed. D. Maximum Intensity projection of confocal images of MDA-MB-231 cells grown in 3D-collagen and transfected with ZsGreen, ZsRASSF1C wild-type or mutant plasmids. Cells were stained with F-actin (red) and pMLCII (far red, false colored to green). Arrows indicate site of cell blebbing. Scale 5 μm. Data are represented as mean ± SEM.* p ≤ 0.05, ** p ≤ 0.01, *** p ≤ 0.001.

**Supplementary Figure 3.**
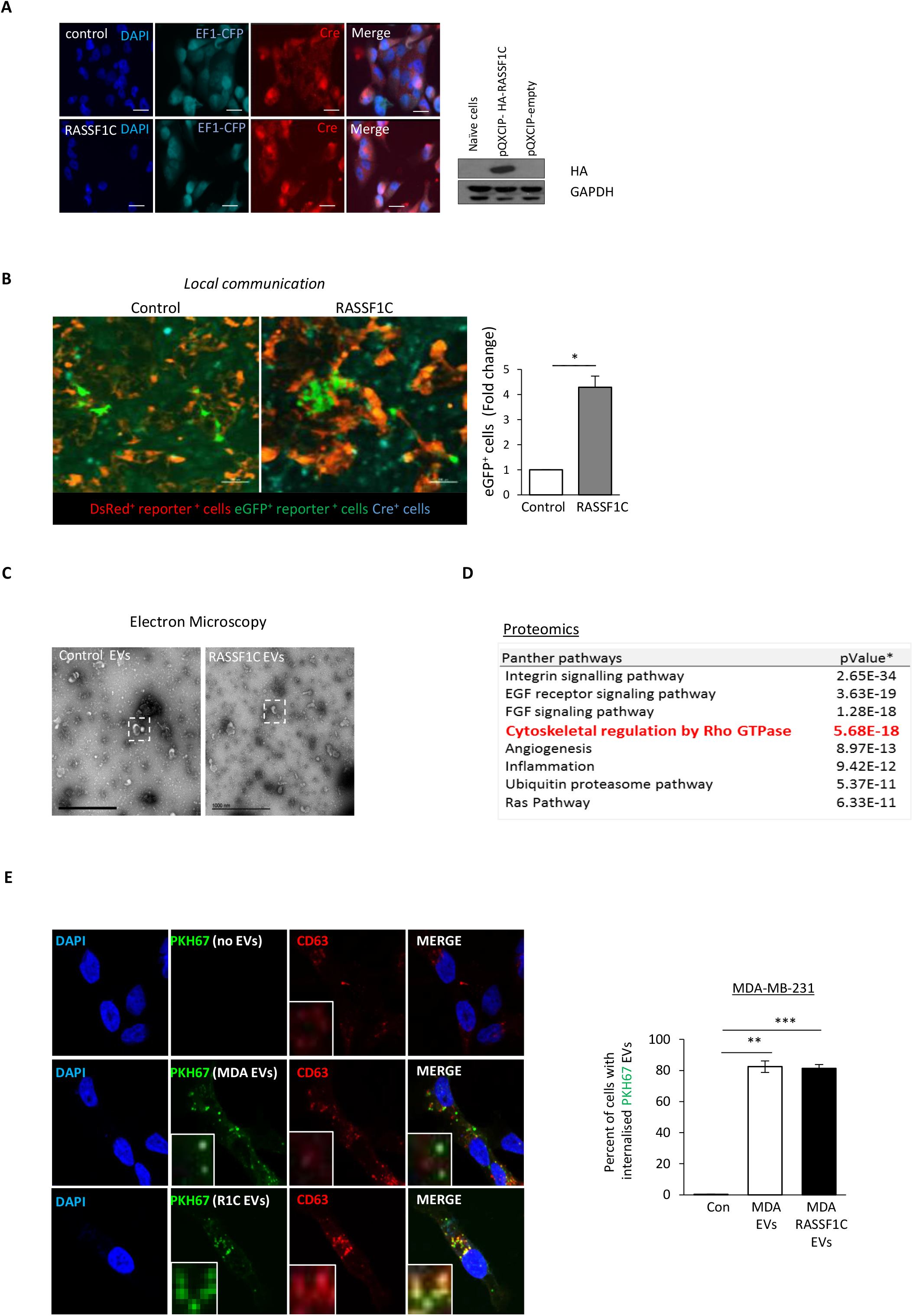
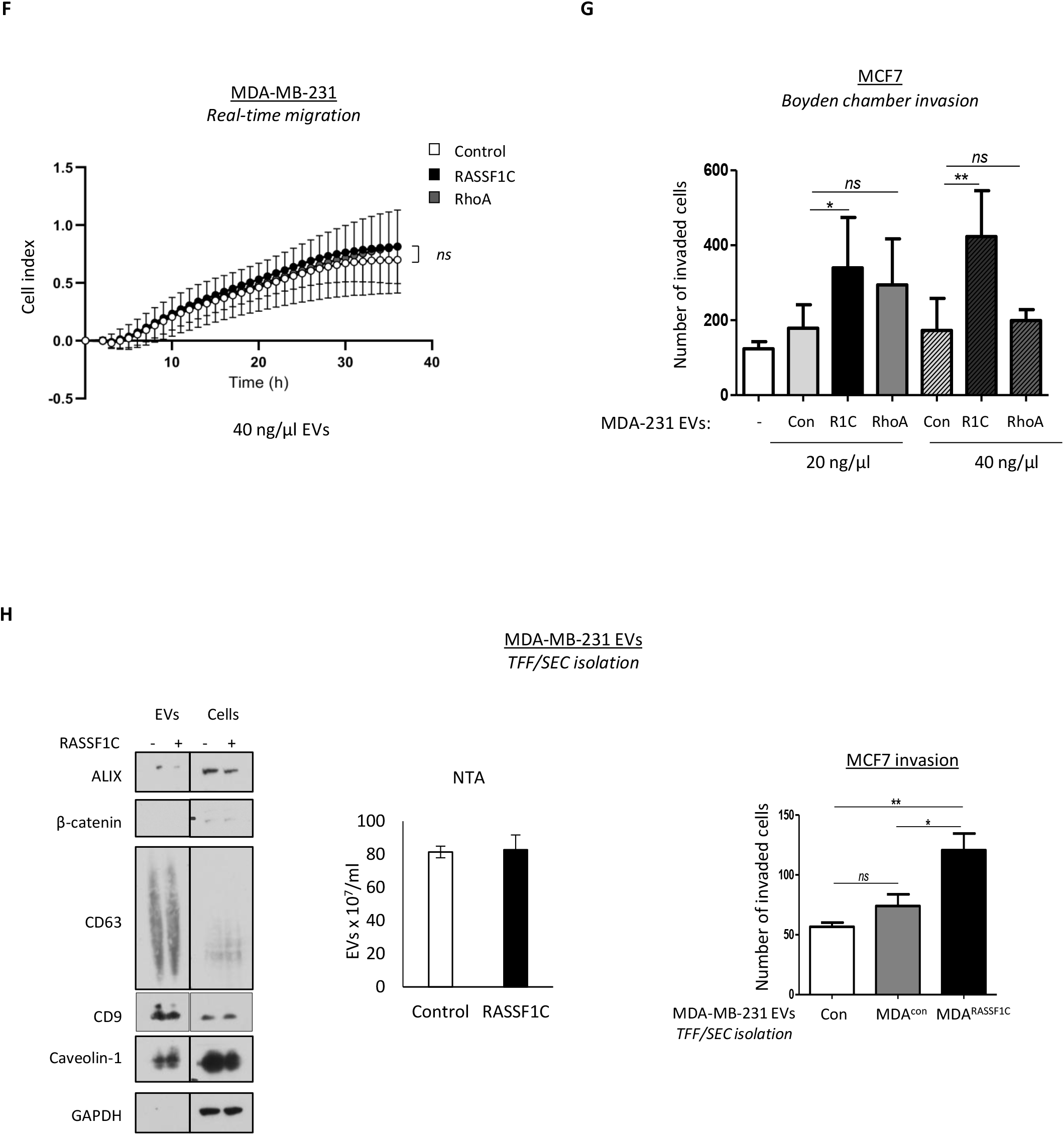
A. Left: Confocal images of MDA-MB-231^CFP;Cre;Control^ or MDA-MB-231^CFP;Cre;HA-RASSF1C^ and stained for Cre expression (red). Scale bars 20 μm. Right: western blots of RASSF1C expression (probed as HA tag) in MDA-MB-231^CFP;Cre;Control^ and MDA-MB-231^CFP;Cre;HA-RASSF1C^. GAPDH was used as loading control. B. Images and quantification of T47D cells that express eGFP^+^ in a mixed culture of T47D reporter cells and MDA-MB-231 donor cells infected with pQXCIP (MDA-MB-231^CFP;Cre;Control^) or pQXCIP-HA-RASSF1C (MDA-MB-231^CFP;Cre;HA-RASSF1C^). Scale bars 100 μm. C. Full view transmission electron microscopy images of EVs isolated from MDA-MB-231 cells expressing Control (pcDNA3) or Flag-RASSF1C, from which magnifications in Fig. 3B are taken. Scale bars 1000 nm. D. Panther pathway analysis of the shared EV proteome of MDA-MB-231 cells transfected with Control (pcDNA3) or FLAG-RASSF1C. E. Representative pictures and quantification of the percentage of MDA-MB-231 cells that have internalized PKH67 stained EVs isolated from MDA-MB-231 expressing Control (MDA EVs) or FLAG-RASSF1C (R1C EVs). Recipient cells have been fixed, stained and imaged 18 h after treatment with 50 ng/μl EVs. F. Related to Fig. 3e, normalized values of real-time invasion rates in xCELLigence plates of MDA-MB-231 cells treated with 40 ng/μl EVs derived from MCF7 cells expressing a Control, FLAG-RASSF1C or EGFP-RhoA plasmid, left to migrate for 36 h. G. Related to Fig. 3f, quantification of Boyden chamber invasion assay in 3D matrigel of MCF7 cells, treated with 20 or 40 ng/μl EVs derived from MDA-MB-231 cells, transfected the same constructs used in Fig. 3e. H. EVs from MDA-MB-231 cells transiently expressing a Control or a Flag-RASSF1C construct were purified by tangential-ultrafiltration (TFF) followed by Size Exclusion Liquid Chromatography (SEC), to ensure purity of the preparations. Left: Western blot analysis of EVs isolated from MDA-MB-231 cells transfected with Control (pcDNA3) or FLAG-RASSF1C with indicated antibodies. Equal amounts of protein were loaded after normalization via microBCA assay, and middle: averaged Nanoparticle Tracking Analysis displaying EV concentration for both conditions. Right: equal amounts of MDA-MB-231 EVs were incubated with MCF7 cells in Boyden chamber invasion assay. Quantification of the number of invaded cells in the presence and absence of EVs is represented. Data are represented as mean ± SEM.* p ≤ 0.05, ** p ≤ 0.01, *** p ≤ 0.001.

**Supplementary Figure 4.**
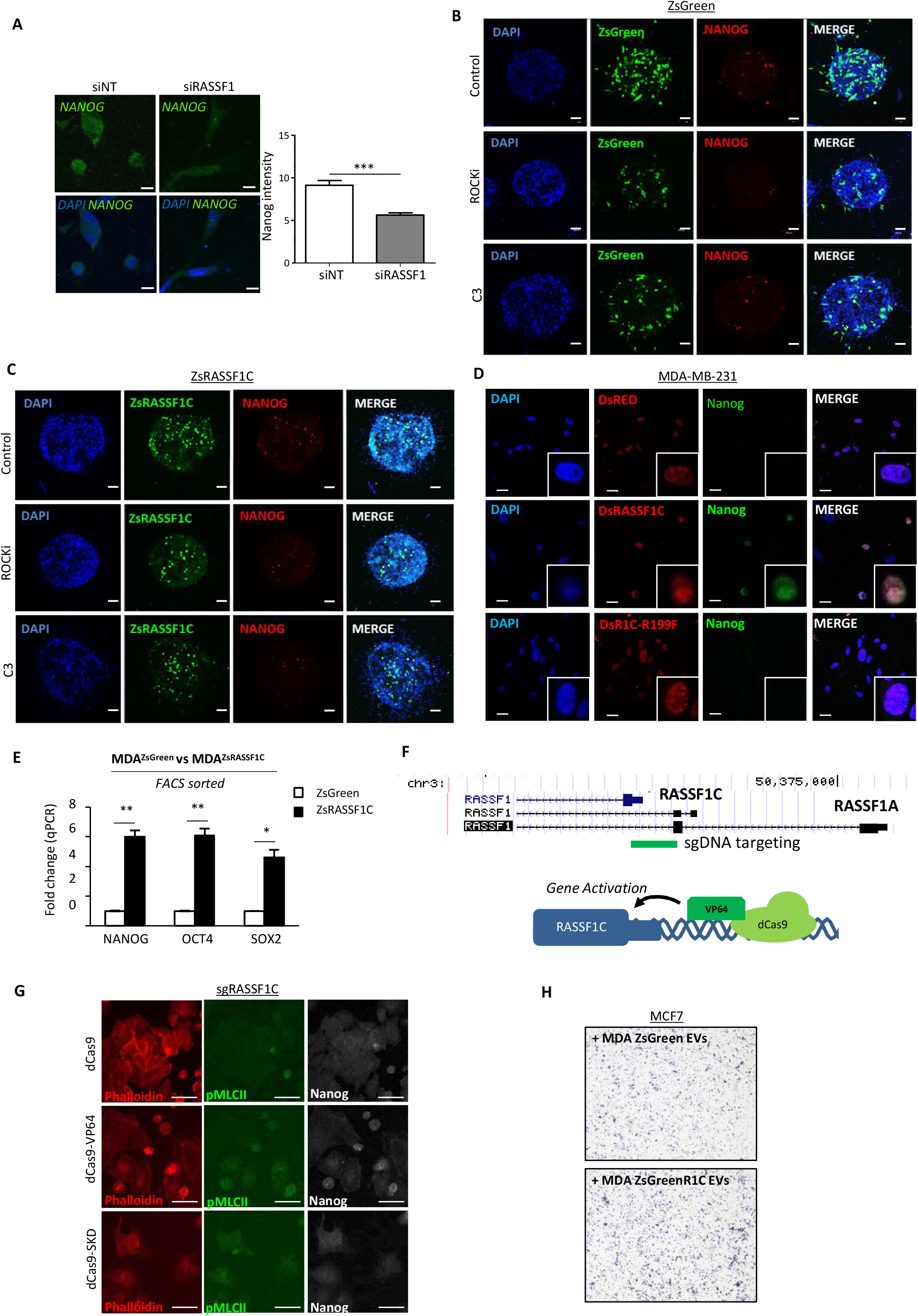
A. Representative confocal images and quantification of fluorescent intensity of Nanog expression in MDA-MB-231 cells after siRNA treatment against RASSF1 (siRASSF1). Scale bars represent 10 μm. B. Confocal images of MDA-MB-231 ZsGreen or C. ZsRASSF1C spheroids treated with ROCK inhibitor (Y-27632, 10 μmol/L) or Rho inhibitor (C3, 2 μg/ml). Cells were stained for Nanog (red). Scale bars represent 200 μm. D. Confocal images of MDA-MB-231 cells grown on 3D-collagen and transfected with Control (DsRed), DsRASSF1C or DsRASSF1C-R199F (DsR1C-R199F). Cells were stained with Nanog (green). Scale bars represent 20 μm. E. Fold change of qRT-PCR of Nanog, OCT4 and SOX2 mRNA from FACS sorted MDA-MB-231 ZsGreen or ZsRASSF1C. F. Cartoon representing the methodology used to induce the expression of RASSF1C using the CRISPR-dCas9 system and genomic location of guide RNA target location. G. Related to Fig. 4h, representative confocal images of MCF7 cells expressing CRISPR-dCas9, dCas9-VP64, dCas9-SKD plus single guide RNA for the RASSF1C promoter (sgRASSF1C) and stained for Phalloidin (red), pMLCII (green) and Nanog (white). Scale bars 20 μm. H. Related to Fig. 4i, representative images of Boyden chamber invasion of MCF7 cells incubated with EVs derived from MDA-MB-231 (control or RASSF1C) cells sorted for Green^high^ expression. Data are represented as mean ± SEM.* p ≤ 0.05, ** p ≤ 0.01, *** p ≤ 0.001.

**Supplementary Figure 5.**
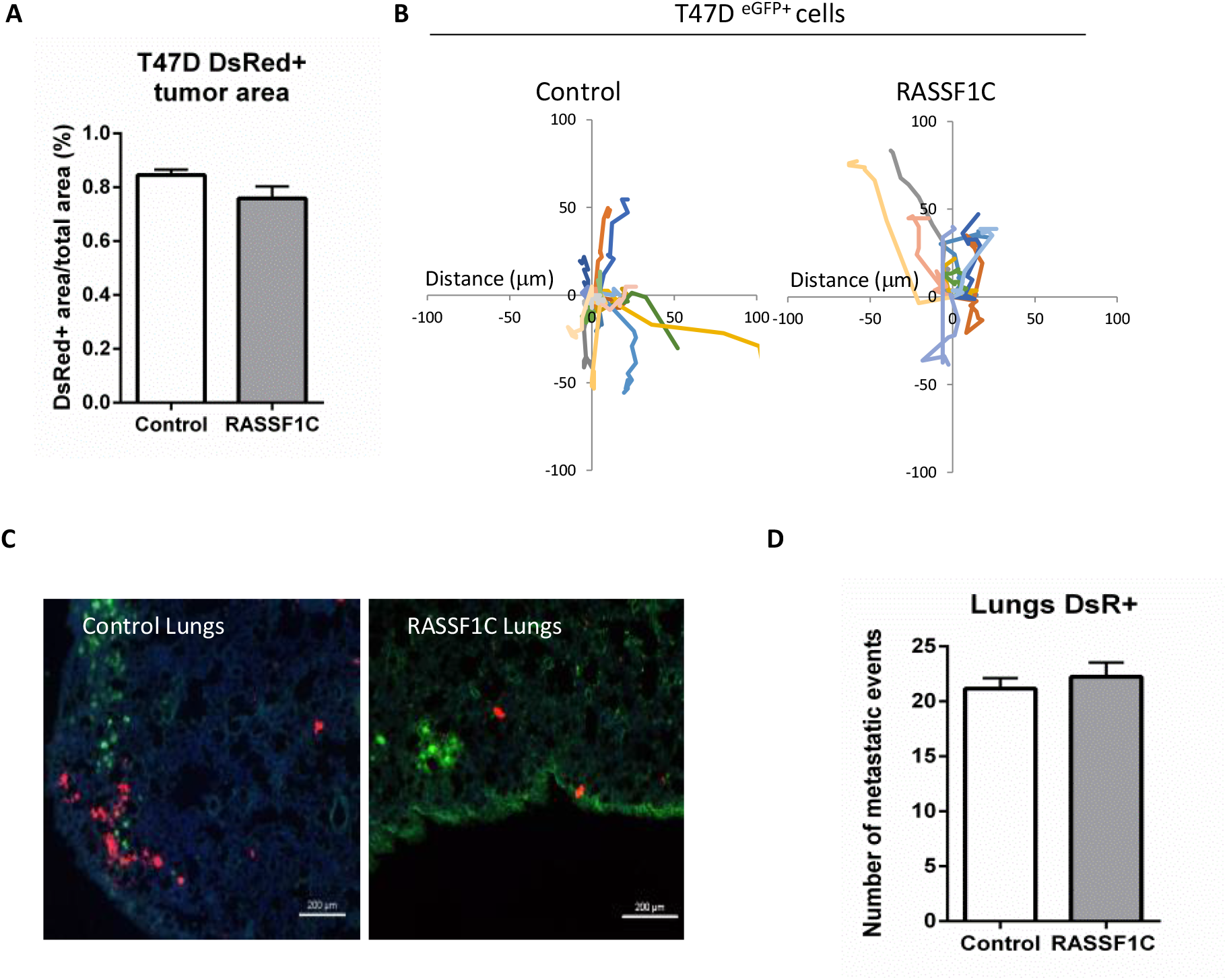
A. Quantification of the size of the T47D^DsRed^ cells within MDA-MB-231^CFP;Cre;Contral^ or MDA-MB-231^CFP;Cre;HA-RASSF1C^ tumors indicating equal contribution to tumor volume. B. Representative 3 hour migration paths of T47D^eGFP^ reporter cells within a single intravital imaging field of tumors co-injected with MDA-MB-231^CFP;Cre;Control^ (Control, left) or MDA-MB-231^CFP;Cre;HA-RASSF1C^ (RASSF1C, right). C. Representative confocal images of lungs of mice with mammary gland tumors initiated by either the coinjection of T47D^DsRed^ reporter cells with MDA-MB-231^CFP;Cre;Contral^ or MDA-MB-231^CFP;Cre;HA-RASSF1C^. Scale bars represent 200 μm. D. Quantification of the number of T47D^DsRed^ metastatic events found in the lungs of mice with mammary gland tumors. Whole lung tile images were used for quantification, 6 of 10 μm sections each, 100 μm apart. Data are represented as mean ± SEM.* p ≤ 0.05, ** p ≤ 0.01, *** p ≤ 0.001.

**Supplementary Figure 6.**
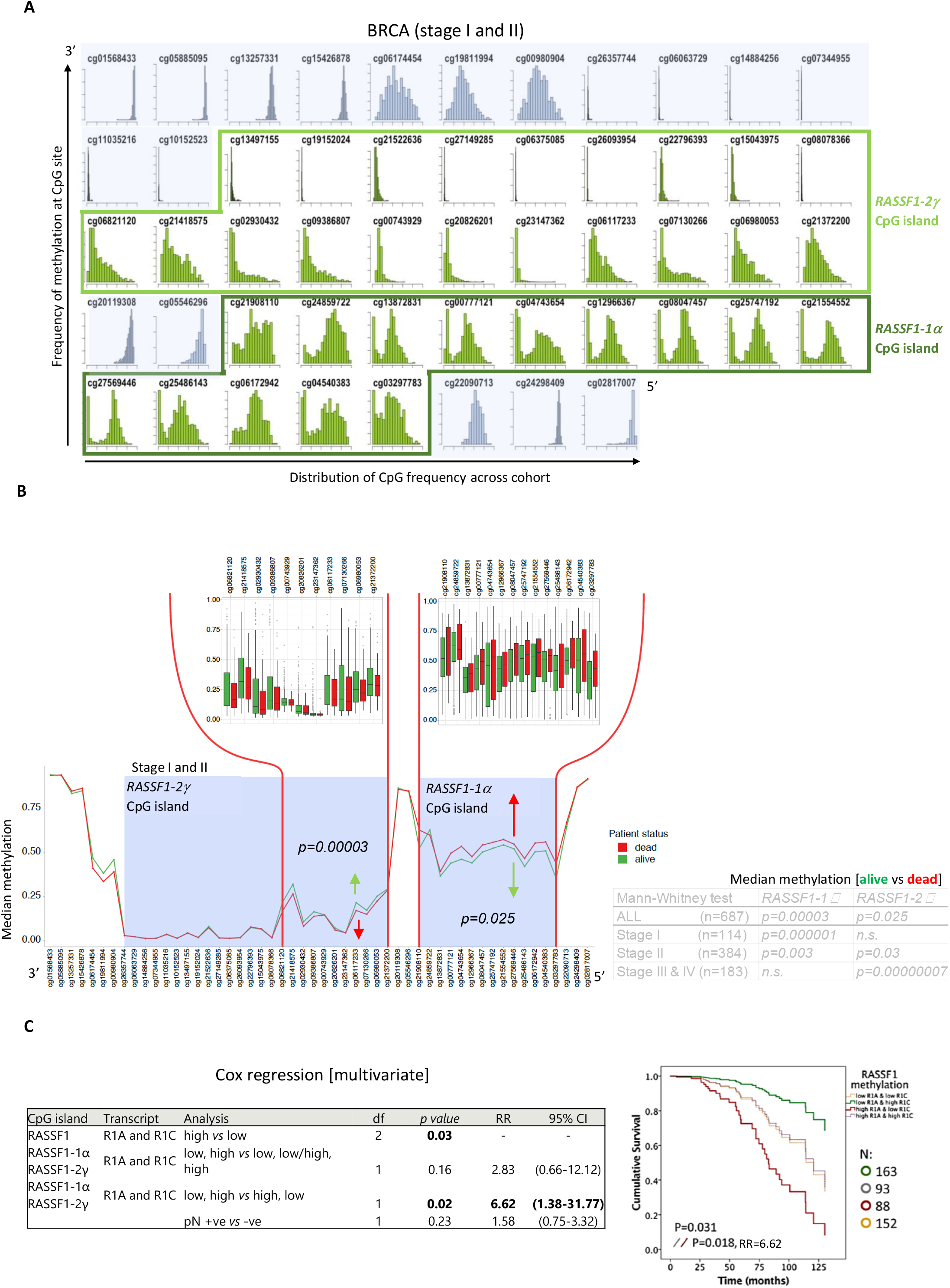
A. Histograms depicting distribution of methylation across stage I and II breast cancer tumours for individual *RASSF1* HM450K CpG sites located within CpG islands (green) or CpG shores (blue). B. Distribution of median methylation of all individual *RASSF1* CpG sites in tumours of stage I and II breast cancer patients who were alive (green) or dead (red) on last follow up. Bars give individual values of the median methylation of patients in the area that gives greatest prognostic value. Table summarizes the differences in median methylation of *RASSF1A* and *RASSF1C* CpGs according to tumor stage, with a particular focus on stage I and II. C. Related to Fig. 6c. Left: table summarizing Cox multivariate analysis performed on the same data-set as in Fig. 6c. Right: Cox survival curves depicting interaction of RASSF1A and RASSF1C methylation to predict the outcome of early stage breast cancer patients. Yellow, low RASSF1A and low RASSF1C; green, low RASSF1A and high RASSF1C; red, high RASSF1A and low RASSF1C; grey, high RASSF1A and high RASSF1C. Abbreviations used in the table: df, degrees of freedom; CI, confidence interval; RR, relative risk; pN, pathologic node status.

**Video 1**

Example of ZsRASSF1C expressing MCF7 cell that became rounded, a morphological change associated with increased contractility. The cells were imaged for up to 8 h at a rate of 1 picture every 15 min.

**Video 2**

Representative intravital imaging video of MDA-MB-231^CFP;Cre;Control^/T47D^DsRed^ tumors where eGFP^+^ recombined reporter cells adopt mesenchymal mode of motility.

**Video 3 and 4**

Representative intravital imaging videos taken from the same tumor at lower (40 μm) and higher (20 μm) magnification of MDA-MB-231^CFP;Cre;HA-RASSF1C^/T47D^DsRed^ tumors where eGFP^+^ recombined reporter cells adopt amoeboid mode of motility.

## References

1. Friedl P, Bröcker EB. The biology of cell locomotion within three-dimensional extracellular matrix. Cell Mol Life Sci. 2000;57(1):41–64. doi:10.1007/s000180050498

2. Friedl P, Locker J, Sahai E, Segall JE. Classifying collective cancer cell invasion. Nat Cell Biol. 2012;(14):777–783. doi:10.1038/ncb2548

3. Álvarez-González B, Meili R, Bastounis E, Firtel RA, Lasheras JC, Del Álamo JC. Three-dimensional balance of cortical tension and axial contractility enables fast amoeboid migration. Biophys J. 2015;108(4):821–832. doi:10.1016/j.bpj.2014.11.3478

4. Kimura K, Ito M, Amano M, et al. Regulation of myosin phosphatase by Rho and Rho-associated kinase (Rho-kinase). Science (80-). 1996;273(5272):245–248. doi:10.1126/science.273.5272.245

5. Egeblad M, Rasch MG, Weaver VM. Dynamic interplay between the collagen scaffold and tumor evolution. Curr Opin Cell Biol. 2010;22(5):697–706. doi:10.1016/j.ceb.2010.08.015

6. He K, Xu T, Goldkorn A. Cancer cells cyclically lose and regain drug-resistant highly tumorigenic features characteristic of a cancer stem-like phenotype. Mol Cancer Ther. 2011;10(6):938–948. doi:10.1158/1535-7163.MCT-10-1120

7. Luga V, Zhang L, Viloria-Petit AM, et al. Exosomes mediate stromal mobilization of autocrine Wnt-PCP signaling in breast cancer cell migration. Cell. 2012;151(7):1542–1556. doi:10.1016/j.cell.2012.11.024

8. Sung BH, Ketova T, Hoshino D, Zijlstra A, Weaver AM. Directional cell movement through tissues is controlled by exosome secretion. Nat Commun. 2015;6(May):7164. http://www.nature.com/doifinder/10.1038/ncomms8164

9. Hoshino A, Costa-Silva B, Shen TL, et al. Tumour exosome integrins determine organotropic metastasis. Nature. 2015;527(7578):329–335. doi:10.1038/nature15756

10. Peinado H, Alečković M, Lavotshkin S, et al. Melanoma exosomes educate bone marrow progenitor cells toward a pro-metastatic phenotype through MET. Nat Med. 2012;18(6):883–891. doi:10.1038/nm.2753

11. Li B, Antonyak MA, Zhang J, Cerione RA. RhoA triggers a specific signaling pathway that generates transforming microvesicles in cancer cells. Oncogene. 2012;31(45):4740–4749. doi:10.1038/onc.2011.636

12. Sedgwick AE, Clancy JW, Olivia Balmert M, D’Souza-Schorey C. Extracellular microvesicles and invadopodia mediate non-overlapping modes of tumor cell invasion. Sci Rep. 2015;5(1):14748. doi:10.1038/srep14748

13. Paluch EK, Raz E. The role and regulation of blebs in cell migration. Curr Opin Cell Biol. 2013;25(5):582–590. doi:10.1016/j.ceb.2013.05.005

14. Lee M-G, Jeong S-I, Ko K-P, et al. RASSF1A Directly Antagonizes RhoA Activity through the Assembly of a Smurf1-Mediated Destruction Complex to Suppress Tumorigenesis. Cancer Res. 2016;76(7):1847–1859. doi:10.1158/0008-5472.CAN-15-1752

15. Pefani DE, Latusek R, Pires I, et al. RASSF1A-LATS1 signalling stabilizes replication forks by restricting CDK2-mediated phosphorylation of BRCA2. Nat Cell Biol. 2014;16(August):962–971, 1-8. doi:10.1038/ncb3035

16. Pefani DE, Pankova D, Abraham AG, et al. TGF-β Targets the Hippo Pathway Scaffold RASSF1A to Facilitate YAP/SMAD2 Nuclear Translocation. Mol Cell. 2016;63(1):156–166. doi:10.1016/j.molcel.2016.05.012

17. Vlahov N, Scrace S, Soto MS, et al. Alternate RASSF1 Transcripts Control SRC Activity, E-Cadherin Contacts, and YAP-Mediated Invasion. Curr Biol. 2015;25(23):3019–3034. doi:10.1016/j.cub.2015.09.072

18. Grawenda AM, O’Neill E. Clinical utility of RASSF1A methylation in human malignancies. Br J Cancer. 2015;113(3):372–381. doi:10.1038/bjc.2015.221

19. Reeves ME, Baldwin SW, Baldwin ML, et al. Ras-association domain family 1C protein promotes breast cancer cell migration and attenuates apoptosis. BMC Cancer. 2010;10:562. doi:10.1186/1471-2407-10-562

20. Canel M, Serrels A, Miller D, et al. Quantitative in vivo imaging of the effects of inhibiting integrin signaling via Src and FAK on cancer cell movement: Effects on e-cadherin dynamics. Cancer Res. 2010;70(22):9413–9422. doi:10.1158/0008-5472.CAN-10-1454

21. DerMardirossian C, Rocklin G, Seo JY, Bokoch GM. Phosphorylation of RhoGDI by Src regulates Rho GTPase binding and cytosol-membrane cycling. Mol Biol Cell. 2006;17(11):4760–4768. doi:10.1091/mbc.E06-06-0533

22. Estrabaud E, Lassot I, Blot G, et al. RASSF1C, an Isoform of the Tumor Suppressor RASSF1A, Promotes the Accumulation of -Catenin by Interacting with TrCP. Cancer Res. 2007;67(3):1054–1061. doi:10.1158/0008-5472.CAN-06-2530

23. Calanca N, Paschoal AP, Munhoz ÉP, et al. The long non-coding RNA ANRASSF1 in the regulation of alternative protein-coding transcripts RASSF1A and RASSF1C in human breast cancer cells: implications to epigenetic therapy. Epigenetics. 2019;14(8):741–750. doi:10.1080/15592294.2019.1615355

24. Chatzifrangkeskou M, Pefani D, Eyres M, et al. RASSF1A is required for the maintenance of nuclear actin levels. EMBO J. 2019;38(16). doi:10.15252/embj.2018101168

25. Montenegro MF, Sáez-Ayala M, Piñero-Madrona A, Cabezas-Herrera J, Rodríguez-López JN. Reactivation of the Tumour Suppressor RASSF1A in Breast Cancer by Simultaneous Targeting of DNA and E2F1 Methylation. PLoS One. 2012;7(12). doi:10.1371/journal.pone.0052231

26. Georgouli M, Herraiz C. Regional Activation of Myosin II in Cancer Cells Drives Tumor Progression via a Secretory Cross-Talk with the Immune Microenvironment. Cell. 2019;176:757–774. doi:10.1016/j.cell.2018.12.038

27. Keller H, Eggli P. Protrusive activity, cytoplasmic compartmentalization, and restriction rings in locomoting blebbing Walker carcinosarcoma cells are related to detachment of cortical actin from the plasma membrane. Cell Motil Cytoskeleton. 1998;41(2):181–193. doi:10.1002/(SICI)1097-0169(1998)41:2<181::AID-CM8>3.0.CO;2-H

28. Wyckoff JB, Pinner SE, Gschmeissner S, Condeelis JS, Sahai E. ROCK- and Myosin-Dependent Matrix Deformation Enables Protease-Independent Tumor-Cell Invasion In Vivo. Curr Biol. 2006;16(15):1515–1523. doi:10.1016/j.cub.2006.05.065

29. Brooks PC, Strömblad S, Sanders LC, et al. Localization of matrix metalloproteinase MMP-2 to the surface of invasive cells by interaction with integrin αvβ3. Cell. 1996;85(5):683–693. doi:10.1016/S0092-8674(00)81235-0

30. Pandya P, Orgaz JL, Sanz-Moreno V. Modes of invasion during tumour dissemination. Mol Oncol. 2017;11(1):5–27. doi:10.1002/1878-0261.12019

31. Sahai E, Marshall CJ. Differing modes for tumour cell invasion have distinct requirements for Rho/ROCK signalling and extracellular proteolysis. Nat Cell Biol. 2003;5(8):711–719. doi:10.1038/ncb1019

32. Snoek-van Beurden PAM, Von Den Hoff JW. Zymographic techniques for the analysis of matrix metalloproteinases and their inhibitors. Biotechniques. 2005;38(1):73–83. doi:10.2144/05381RV01

33. Sahai E, Marshall CJ. Differing modes of tumour cell invasion have distinct requirements for Rho/ROCK signalling and extracellular proteolysis. Nat Cell Biol. 2003;5(8):711–719. doi:10.1038/ncb1019

34. Zomer A, Maynard C, Verweij FJ, et al. In Vivo Imaging Reveals Extracellular Vesicle-Mediated Phenocopying of Metastatic Behavior. Cell. 2015;161(5):1046–1057. doi:10.1016/j.cell.2015.04.042

35. Friedl P, Zänker KS, Bröcker EB. Cell migration strategies in 3-D extracellular matrix: Differences in morphology, cell matrix interactions, and integrin function. Microsc Res Tech. 1998;43(5):369–378. doi:10.1002/(SICI)1097-0029(19981201)43:5<369::AID-JEMT3>3.0.CO;2-6

36. Palecek SP, Loftust JC, Ginsberg MH, Lauffenburger DA, Horwitz AF. Integrin-ligand binding properties govern cell migration speed through cell-substratum adhesiveness. Nature. 1997;385(6616):537–540. doi:10.1038/385537a0

37. Quail DF, Joyce JA. Microenvironmental regulation of tumor progression and metastasis. Nat Med. 2013;19(11):1423–1437. doi:10.1038/nm.3394

38. Willms E, Johansson HJ, Mäger I, et al. Cells release subpopulations of exosomes with distinct molecular and biological properties. Sci Rep. 2016;6(1):22519. doi:10.1038/srep22519

39. Willms E, Cabañas C, Mäger I, Wood MJA, Vader P. Extracellular vesicle heterogeneity: Subpopulations, isolation techniques, and diverse functions in cancer progression. Front Immunol. 2018;9(APR). doi:10.3389/fimmu.2018.00738

40. Pietrovito L, Leo A, Gori V, et al. Bone marrow-derived mesenchymal stem cells promote invasiveness and transendothelial migration of osteosarcoma cells via a mesenchymal to amoeboid transition. Mol Oncol. 2018;12(5):659–676. doi:10.1002/1878-0261.12189

41. Taddei M, Giannoni E, Morandi A, et al. Mesenchymal to amoeboid transition is associated with stemlike features of melanoma cells. Cell Commun Signal. 2014;12(1):24. doi:10.1186/1478-811X-12-24

42. Ginestier C, Hur MH, Charafe-Jauffret E, et al. ALDH1 Is a Marker of Normal and Malignant Human Mammary Stem Cells and a Predictor of Poor Clinical Outcome. Cell Stem Cell. 2007;1(5):555–567. doi:10.1016/j.stem.2007.08.014

43. Ponti D, Costa A, Zaffaroni N, et al. Isolation and in vitro propagation of tumorigenic breast cancer cells with stem/progenitor cell properties. Cancer Res. 2005;65(13):5506–5511. doi:10.1158/0008-5472.CAN-05-0626

44. Rasti A, Mehrazma M, Madjd Z, Abolhasani M, Saeednejad Zanjani L, Asgari M. Co-expression of Cancer Stem Cell Markers OCT4 and NANOG Predicts Poor Prognosis in Renal Cell Carcinomas. Sci Rep. 2018;8(1):1–11. doi:10.1038/s41598-018-30168-4

45. Ramos EAS, Camargo AA, Braun K, et al. Simultaneous CXCL12 and ESR1 CpG island hypermethylation correlates with poor prognosis in sporadic breast cancer. BMC Cancer. 2010;10(1):23. doi:10.1186/1471-2407-10-23

46. Malpeli G, Amato E, Dandrea M, et al. Methylation-associated down-regulation of RASSF1A and up-regulation of RASSF1Cin pancreatic endocrine tumors. BMC Cancer. 2011;11(1):351. doi:10.1186/1471-2407-11-351

47. Yamashita K, Hosoda K, Nishizawa N, Katoh H, Watanabe M. Epigenetic biomarkers of promoter DNA methylation in the new era of cancer treatment. Cancer Sci. 2018;109(12):3695–3706. doi:10.1111/cas.13812

48. Quail DF, Joyce JA. Microenvironmental regulation of tumor progression and metastasis. Nat Med. 2013;19(11):1423–1437. doi:10.1038/nm.3394

49. Wortzel I, Dror S, Kenific CM, Lyden D. Exosome-Mediated Metastasis: Communication from a Distance. Dev Cell. 2019;49(3):347–360. doi:10.1016/j.devcel.2019.04.011

50. Van Niel G, D’Angelo G, Raposo G. Shedding light on the cell biology of extracellular vesicles. Nat Rev Mol Cell Biol. 2018;19(4):213–228. doi:10.1038/nrm.2017.125

51. Harris D a., Patel SH, Gucek M, Hendrix A, Westbroek W, Taraska JW. Exosomes Released from Breast Cancer Carcinomas Stimulate Cell Movement. PLoS One. 2015;10(3):e0117495. doi:10.1371/journal.pone.0117495

52. Costa-Silva B, Aiello NM, Ocean AJ, et al. Pancreatic cancer exosomes initiate pre-metastatic niche formation in the liver. Nat Cell Biol. 2015;17(6):816–826. doi:10.1038/ncb3169

53. Peinado H, Zhang H, Matei IR, et al. Pre-metastatic niches: Organ-specific homes for metastases. Nat Rev Cancer. 2017;17(5). doi:10.1038/nrc.2017.6

54. Volodko N, Salla M, Zare A, et al. RASSF1A Site-Specific Methylation Hotspots in Cancer and Correlation with RASSF1C and MOAP-1. Cancers (Basel). 2016;8(6):55. doi:10.3390/cancers8060055

55. Lawson DA, Bhakta NR, Kessenbrock K, et al. Single-cell analysis reveals a stem-cell program in human metastatic breast cancer cells. Nature. 2015;526(7571):131–135. Accessed August 3, 2017. http://www.ncbi.nlm.nih.gov/pubmed/26416748

56. Papaspyropoulos A, Bradley L, Thapa A, et al. RASSF1A uncouples Wnt from Hippo signalling and promotes YAP mediated differentiation via p73. Nat Commun. 2018;9(1):424. doi:10.1038/s41467-017-02786-5

57. Pankova D, Jiang Y, Chatzifrangkeskou M, et al. RASSF 1A controls tissue stiffness and cancer stem-like cells in lung adenocarcinoma. EMBO J. 2019;38(13):e100532. doi:10.15252/embj.2018100532

58. Dupont S, Morsut L, Aragona M, et al. Role of YAP/TAZ in mechanotransduction. Nature. 2011;474(7350):179–183. doi:10.1038/nature10137

59. Cordenonsi M, Zanconato F, Azzolin L, et al. The hippo transducer TAZ confers cancer stem cell-related traits on breast cancer cells. Cell. 2011;147(4):759–772. doi:10.1016/j.cell.2011.09.048

60. Foty R. A simple hanging drop cell culture protocol for generation of 3D spheroids. J Vis Exp. 2011;(51). doi:10.3791/2720

61. Pozzi A, Moberg PE, Miles LA, Wagner S, Soloway P, Gardner HA. Elevated matrix metalloprotease and angiostatin levels in integrin α1 knockout mice cause reduced tumor vascularization. Proc Natl Acad Sci U S A. 2000;97(5):2202–2207. doi:10.1073/pnas.040378497

62. Cano-Rodriguez D, Gjaltema RAF, Jilderda LJ, et al. Writing of H3K4Me3 overcomes epigenetic silencing in a sustained but context-dependent manner. Nat Commun. 2016;7. doi:10.1038/ncomms12284

